# Multiscale brain-wide mapping of α-synuclein-driven dopaminergic degeneration and white matter impairment in α-synucleinopathy mice

**DOI:** 10.64898/2025.12.05.692595

**Authors:** Benjamin F. Combes, Sihan Dong, Martin T. Henrich, Alicia K. Rosenau, Maria Karatsoli, Jie Lu, Sepp Kollmorgen, Marco Reisert, Xunbin Wei, Tian Zeng, Hongjiang Wei, Yi Chen, Liqin Yang, Kuangyu Shi, Peter R Nilsson, Theofanis Karayannis, Rong Chen, Valina L. Dawson, Ted M. Dawson, Wolfgang H. Oertel, Christoph Hock, Daniel Razansky, Axel Rominger, Roger M. Nitsch, Fanni F. Geibl, Ruiqing Ni

## Abstract

The spatiotemporal relationships among α-synuclein inclusion formation, dopaminergic degeneration, and white matter microstructural changes remain poorly defined. To address this, we unilaterally injected preformed fibrils (PFFs) or monomeric α-synuclein into the substantia nigra pars compacta (SNc) of wild-type mice. At 12 and 20 weeks post-injection (wpi), we performed ex vivo magnetic resonance imaging (MRI) at 9.4T to assess microstructural, volumetric and paramagnetic changes, along with light-sheet microscopy (LSM) to map α-synuclein aggregates, and dopaminergic neuron at single-cell resolution across the whole brain. We developed a Python-based, automated registration pipeline that achieves <40 µm alignment error between ex vivo MRI and LSM and computes clearing-induced distortion, enabling scalable deformation analysis and multimodal multiscale data integration. Unilateral SNc injection of α-syn PFFs induced dense phospho-α-syn pathology in the ipsilateral SNc at 12 wpi, which decreased by 20 wpi in parallel with the loss of tyrosine hydroxylase-positive dopaminergic neuron. Pathogenic α-synuclein spreads along with dopaminergic denervation in the nigrostriatal pathway to the ipsilateral striatum and contralateral SNc. Our ex vivo MRI-LSM platform provides a scalable, open-source framework for understanding circuit-level and whole-brain propagation of pathology in an α-synucleinopathy model and beyond.

**Highlights:**

- Open-source, automated pipeline aligns ex vivo 9.4T MRI with light-sheet microscopy (LSM) at <40 µm error, quantifies clearing-induced tissue distortion, and enables multiscale, whole-brain multimodal analysis.
- Whole-brain, single-cell-resolution mapping of α-synuclein pathology and dopaminergic neuron loss in an α-synucleinopathy mouse model using an optimized iDISCO+ protocol that achieves uniform antibody penetration even in dense regions like the striatum.
- Unilateral SNc injection of α-syn PFFs drives spread of p-α-syn pathology along the nigrostriatal pathway to the ipsilateral striatum, BST, CEA, and contralateral SNc, along with dopaminergic degeneration.
- Ipsilateral SNc shows high pS129 pathology at 12 weeks post-injection but significant reduction by 20 wpi, concurrent with loss of tyrosine hydroxylase-positive neurons.
- Multiparametric MRI reveals microstructural and iron-related alterations in striatum, SNc, and VTA, with cross-modal correlations linking pS129 burden, TH loss, and magnetic susceptibility changes.

## Introduction

Parkinson’s disease (PD) is the second most common neurodegenerative disease, affecting more than 10 million people worldwide. The key neuropathological feature of PD is the intracellular accumulation of Lewy bodies (LB) composed of misfolded insoluble alpha-synuclein (α-syn) fibrils that propagate between cells and induce the aggregation of endogenous proteins in recipient cells, leading to LB formation ^1–3^. The formation of LBs in the dopaminergic neurons of the substantia nigra pars compacta (SNc), which are responsible for the degeneration of the nigrostriatal pathway and largely contribute to the onset of characteristic motor symptoms that include bradykinesia, resting tremor, and/or rigidity^4,5^. Advancements in neuroimaging have improved the characterization of PD and have moved toward biomarker-based diagnosis ^6,7^, such as for identifying dopaminergic dysfunction by DaTSCAN using single-photon emission computed tomography. Imaging of α-syn pathologies by using positron emission tomography (PET) has not yet been successful in patients with PD. White matter microstructural alterations detected by diffusion tensor imaging (DTI) ^8^ and iron accumulation detected by quantitative susceptibility mapping (QSM) magnetic resonance imaging (MRI)^9^ have been reported in patients with PD. Mouse models recapitulating PD and α-synucleinopathies have enabled a better understanding of disease mechanisms and the development of biomarkers and treatment strategies.

The absence of disease-modifying therapies and the challenge of early diagnosis emphasize the need for a deeper understanding of the mechanisms driving the spread of α-syn and its connection to neurodegeneration and connectivity alterations in vulnerable circuits. α-Syn preformed-fibrils (PFFs) mice as well as transgenic models have been developed to study the propagation and links among protein aggregation, neuroinflammation, and neurodegeneration ^10–16^. Currently, α-syn distribution pattern evaluation can be performed *ex vivo* by classic immunolabeling methods in animal models, which is tedious. In vivo PET and large-field fluorescence imaging in animal models provide limited resolution, especially for α-syn aggregates of micronmeter size ^17,18^, whereas two-photon imaging using genetically encoded fluorescent reporters to visualize α-syn pathology offers a limited field of view in which reaching subcortical regions that are important for PD is difficult^19^.

Elucidating the propagation of abnormal α-syn inclusions throughout the brain and their spatiotemporal relationship with neurodegeneration and microstructural changes remains a crucial yet unresolved challenge. Whether α-syn levels increase or decrease with the loss of dopaminergic neurons in PD patients during disease progression has not been determined. Here we hypothesize that α-syn pathology leads to dopaminergic denervation, microstructural and iron-related changes, and that the α-syn pathology reduces in early affected regions with the loss of dopaminergic neuron at later stage. Addressing this requires the development of platforms that enable the efficient integrated study of pathological processes across different scales. MRI achieves whole-brain coverage and excellent anatomical contrast, making this technique capable of characterizing gross morphological changes during disease progression, as shown by the detection of cerebral microstructural changes in PD rodent models. Recent advances, particularly organ clearing and light-sheet microscopy (LSM)^20–22^, have enabled comprehensive mapping of neuronal systems, as well as systemic studies at cellular resolution in disease models, in disease models. LSM for amyloid-beta and tau in the mouse and human brains ^23,24^ has been established; however, a previous LSM study of α-syn revealed limited antibody penetration into dense striatal region, which is critical in PD^25^.

Integrating LSM with MRI enables comprehensive, multiscale characterization of the mouse brain. However, accurate quantitative cross-modal registration remains challenging because of distinct contrast mechanisms, illumination nonuniformities (e.g., striping and attenuation), and significant nonlinear deformations introduced by tissue clearing. Standard affine transformations or direct registration to brain atlases often fail to capture severe anisotropic tissue shrinkage and local distortions. Here, we developed an automated, open-source pipeline implemented in Python that encompasses preprocessing and bidirectional registration and computes clearing-induced distortion. This pipeline enables unbiased computational analysis of MRI and LSM data from mouse brains. Using this pipeline, we demonstrated high-resolution whole-brain mapping of α-syn spreading from the SNc and dopaminergic dysfunction by using an LSM (at single-cell resolution, with an adapted iDISCO+ protocol) and microstructural and iron-related changes alterations by using MRI in mouse models.

## Results

### Multiscale imaging and analysis pipeline in the α-syn PFF SNc injection model

The workflow included 9.4 T ex vivo multiparametric MRI and LSM (MesoSPIM) in mice at 12 and 20 weeks following unilateral α-syn PFF or monomer injection into the SNc (**SFig. 1**). Multimodal automated coregistration was performed to map and align pS129 α-syn and dopaminergic impairments with MRI at the whole-brain level. An automated multimodal computational pipeline encompassing multiatlas ROI segmentation, threshold-based pathology extraction, multiscale LSM/MRI fusion, and inter-ROI correlation coefficient network construction was constructed (**Fig 1**).

**Figure 1.**
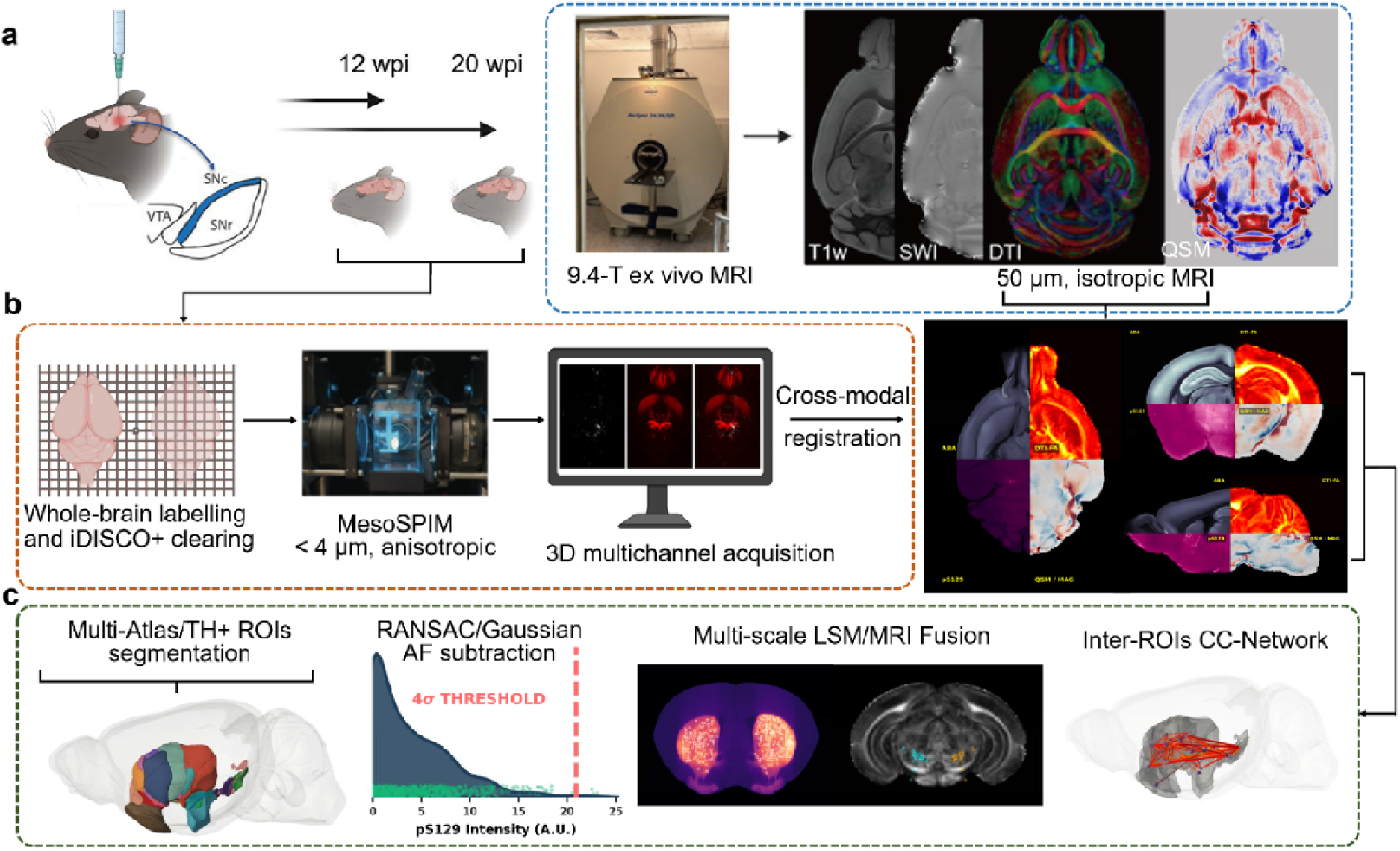
Experimental workflow and multimodal integration pipeline. **(a)** Experimental workflow and 9.4 T ex vivo multiparametric MRI acquisition in mice sacrificed at 12 and 20 weeks following unilateral α-synuclein PFFs or monomer injection into the substantia nigra pars compacta (SNc). (**b)** An optimized iDISCO+ clearing protocol, MesoSPIM imaging (resolution < 4 μm), and multi-modal co-registration was performed to map and align pS129 α-syn and dopaminergic impairments with MRI at the whole-brain level. **(c)** Automated multi-modal computational pipeline encompassing multi-atlas ROI segmentation, threshold-based pathology extraction, multi-scale LSM/MRI fusion, and inter-ROI CC-network construction. CC-Network: correlation coefficient network, PFFs: preformed fibrils, pS129: phosphorylated serine of α-syn at the position 129, SNc: substantia nigra pars compacta, TH: tyrosine hydroxylase. The figure was created with BioRender and Python.

We first histologically validated the α-syn PFF SNc injection and α-syn pattern. At 12 wpi, the mice were sacrificed, and their brains were examined using classical immunohistochemistry against phosphorylated α-syn. Dense α-syn pS129-positive pathology was observed in the TH+ dopaminergic nigral neurons of α-syn PFF-injected mice (**SFig. 2**). β-sheet-detecting compounds AOI987 and hFTAA^26,27^ robustly labeled the intracellular α-syn pS129-positive aggregates (**SFig. 2**), characterized the fibrillar morphology, and allowed us to distinguish between aggregates located in the soma and those in neurites (**SFig. 2**). Furthermore, we observed that α-syn pathology had spread unilaterally to nearby brain regions, such as the ventral segmental area (VTA). It also reached more distant rostral areas, including the striatum, temporal cortex, nucleus accumbens (NAc), central amygdalar nucleus (CEA), and caudal regions, such as the periaqueductal gray (PAG) and pedunculopontine nucleus (PPN) (**Fig. 2g**). We also validated the results using another anti-pS129 α-syn antibody clone (81A) in addition to EP1536Y. Both antibodies target the same pathological form of α-syn and reveal comparable soma and neuritic inclusions in our model (**SFig. 3**). Only sparse α-syn pathology was detected in the contralateral hemisphere, except for caudal regions (**Fig. 2g**). The injection of α-syn monomers did not induce any seeding of pS129 α-syn pathology.

**Figure 2.**
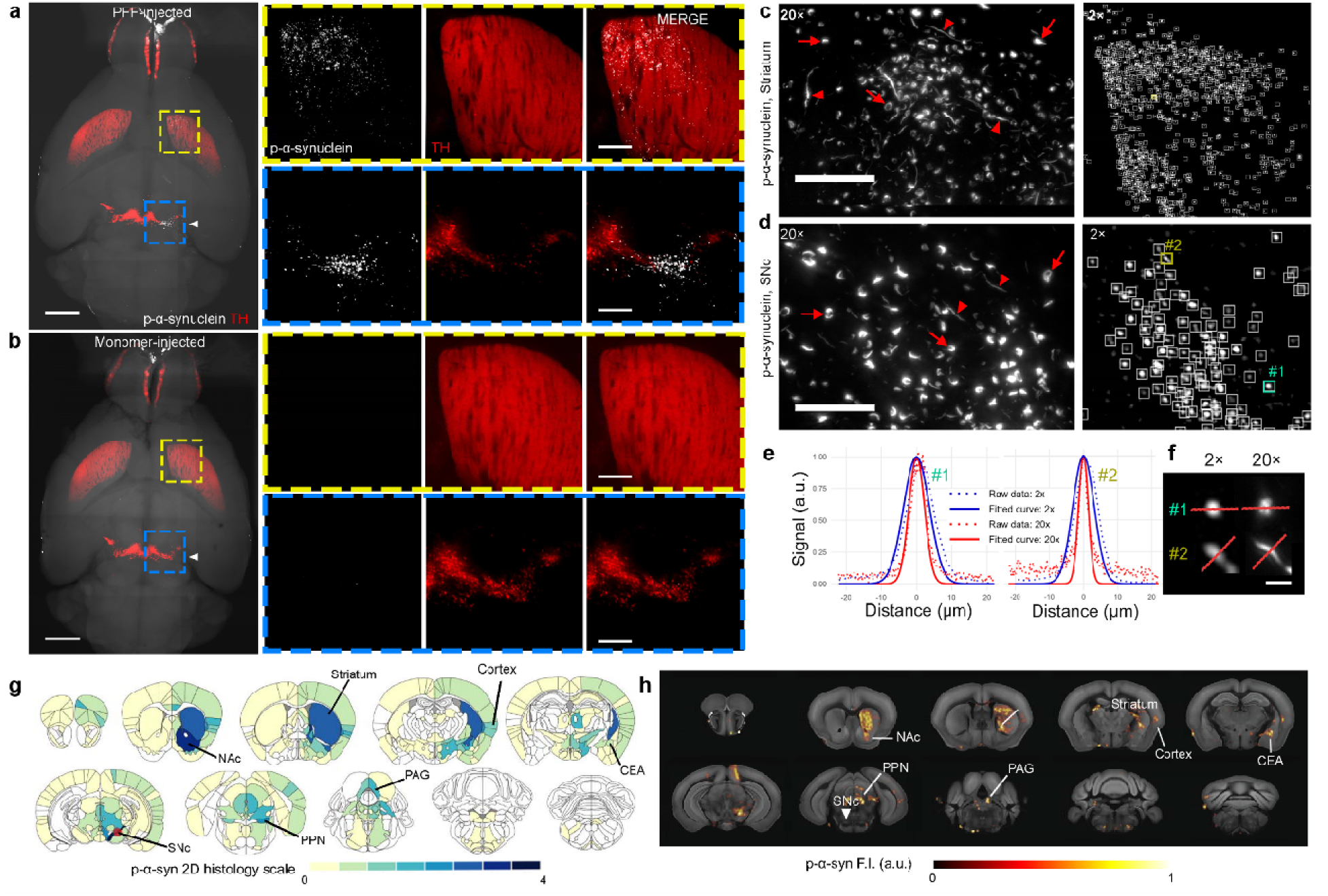
High-resolution whole-brain mapping of α-syn aggregates in PFF-injected mice using an optimized iDISCO+ protocol. **(a)** and **(b)** Representative horizontal slices of whole-brain iDISCO+ staining showing robust and consistent pS129 α-syn (white) and TH (red) labelling in the striatum (yellow boxes) and the SNc (blue boxes) 12 wpi of α-syn PFFs in the SNc (white triangle), with no detectable α-syn signal in monomer-injected controls (b); scale bars = 1.5 mm (overview) and 400 µm (digital zoom, post-capture enlargement). **(c)** Left: 100 µm-thick MIP of higher magnification (20×) from whole-cleared brains reveals the characteristic morphology of α-syn aggregates and confirms the staining specificity in the striatum. Red arrows show soma pathology, while red triangles indicate neurite pathology; scale bar = 100 µm. Right: Representative slices of the striatum show single α-syn aggregates detection using a customed automatic pipeline at 12 wpi; scale bar = 500 µm. **(d)** Left: 100 µm-thick MIP of higher magnification (20×) from whole-cleared brains reveals the characteristic morphology of α-syn aggregates and confirms the staining specificity in the SNc. Red arrows show soma pathology, while red triangles indicate neurite pathology; scale bar = 100 µm. Right: Representative slices of the SNc show single α-syn aggregates detection using a customed automatic pipeline at 12 wpi; scale bar = 100 µm. Green (1) and yellow (2) boxes show the capacity to detect the same single aggregates within the intact cleared brain, and are, in turn, automatically detected. **(e)** The signal profiles of these aggregates obtained at 2× and 20× magnifications are compared along the corresponding red lines. **(f)** Aggregate #1 represents somatic pathology, while aggregate #2 corresponds to neuritic pathology. The Gaussian-fitted full width at half maximum (FWHM) was calculated using the plot profile function in Fiji by plotting the fitting Gaussian curve for aggregate #1 is 8.6 µm at 2× and 5.2 µm at 20×, and for aggregate #2 is 7.3 µm at 2× and 3.3 µm at 20×; scale bar = 10 µm. **(g)** Semiquantitative heatmap depicting the brain-wide α-syn pathology 12 weeks after PFFs injection into the SNc of WT mice. Red dot indicates the injection site. Classical immunohistochemistry and LSM were performed on separate cohorts of mice. **(h)** 3D quantified heatmap in one brain depicting the brain-wide α-syn pathology 12 weeks after PFFs injection into the SNc of WT mice. White triangle indicates the injection site. α-syn: α-synuclein, CEA: central amygdalar nucleus, FWHM: full width at half maximum, hFTAA: heptamer formyl thiophene acetic acid, LSM: light-sheet microscopy, MIP: maximum intensity projection, NAc: nucleus accumbens, PAG: periaqueductal gray, PFFs: preformed fibrils, PPN: pedunculopontine nucleus, pS129: phosphorylated serine of α-syn at the position 129, SNc: substantia nigra pars compacta, TH: tyrosine hydroxylase, wpi: weeks post injection, WT: wild-type.

**Figure 3.**
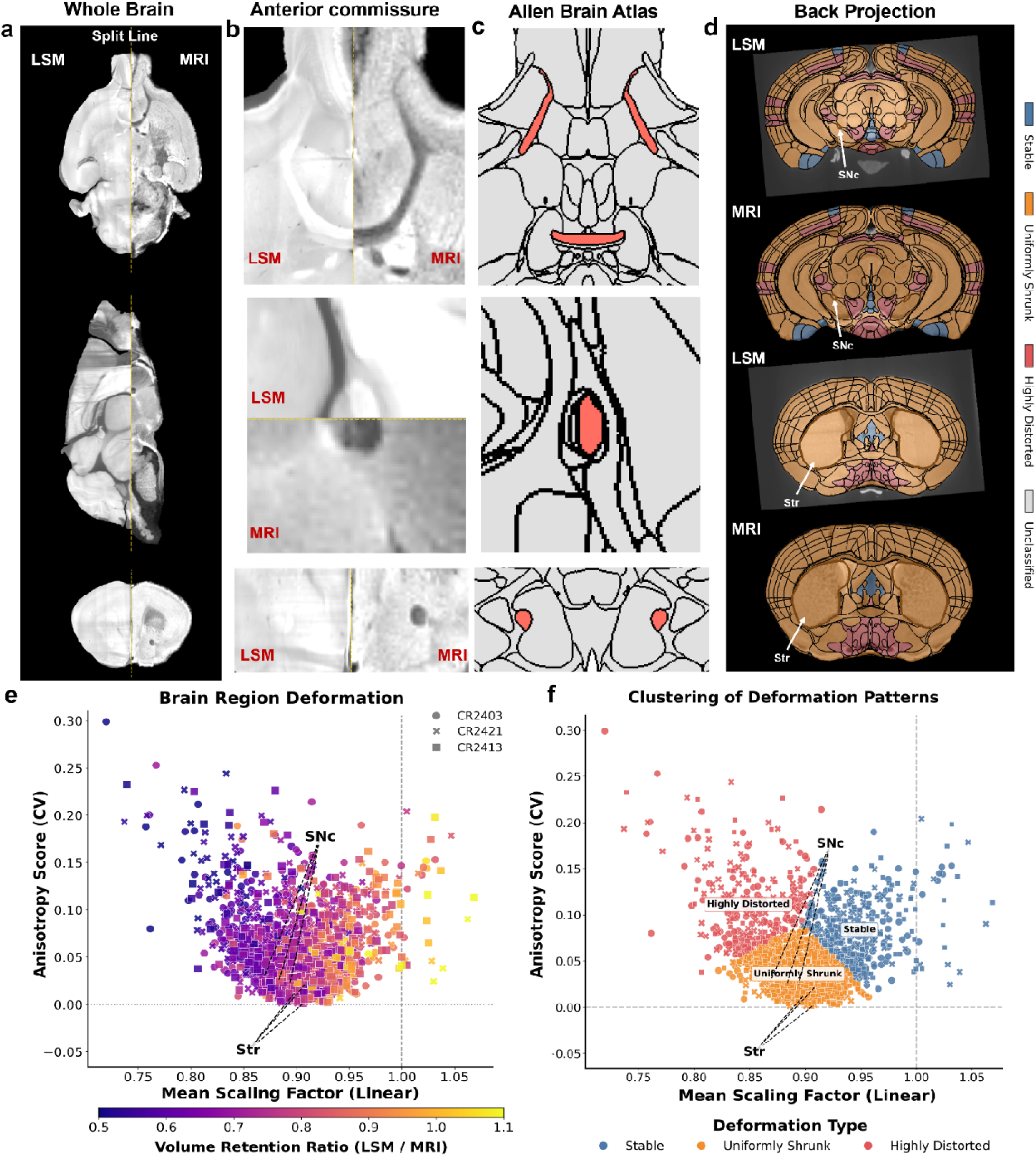
Quantitative evaluation of co-registration accuracy and clearing-induced distortion. **(a-c)** Landmark-based validation of the anatomical alignment of the anterior commissure (olfactory limb) between LSM and MRI displayed at (**a**) the whole brain level, (**b**) a magnified view of the landmark region, and (c) the corresponding ABA reference region. Images are presented in horizontal, sagittal, and coronal planes (from top to bottom). **(d)** Inverse mapping to project ABA annotations into native LSM and native MRI spaces by reversing the registration transformation chain. The colormap representing the distortion patterns is derived from the analyses in (**e**) and (**f**). **(e)** Assessment of morphological distortion via PCA-derived equivalent ellipsoids for regions of interest. **(f)** K-means clustering (k=3) of deformation profiles based on the standardized mean scaling factor and anisotropy score. Regions were classified into three categories characterized by shape fidelity and anisotropy: stable (18.1%), uniformly shrunk (67.8%), and highly distorted (14.1%). ABA: Allen Brain Atlas, LSM: Light-sheet microscopy, MRI: Magnetic resonance imaging, PCA: Principal component analysis.

### The adapted iDISCO+ protocol enables high-resolution, whole-brain detection of individual α-syn pathology and TH-positive neurons

We evaluated the original iDISCO+ protocol and performed optimization. The incubation of the primary and secondary antibodies for 7 days each at 37°C resulted in labeling across many brain regions, including the SNc, VTA, retrorubral field (RRF), cortex, CEA, PAG and PPN (**SFig 4, SVideo 1**). However, in the striatum, only the outer portion exhibited TH labeling, and no pS129 signal was detected, which was expected according to conventional histology (**Fig. 2c, SVideo 1**). We confirmed that this was due to limited antibody penetration with the original iDISCO+ protocol by sectioning the PFF-injected cleared brain into 2 mm-thick coronal slices and rehydrating and relabeling them before clearing and imaging, so-called “reverse” iDISCO+. Following this approach, dense α-syn pS129-positive pathology and TH-positive signals were observed throughout the striatum as well as in the PAG (**SFig. 3**).

Therefore, we modified the protocol to overcome the limited antibody penetration issue^10^. We employed an anti-pS129 α-syn antibody directly conjugated to Alexa555 together with conjugated TH-VioR667 and extended the incubation period to 4 weeks at 37°C. This approach resulted in strong and dense α-syn pS129-positive labeling in the expected regions, including the striatum, with homogeneous TH staining (**Fig. 2a**, **SVideo 2**). High-resolution whole-brain imaging (2.75×2.75×4 µm) acquired at 2× magnification further revealed minimal ring-shaped background at the brain surface (which is characteristic of poor antibody penetration). This enabled the 3D visualization of both α-syn aggregate spread and TH expression in the mouse brain (**SVideo. 3**). The selectivity of the pS129 α-syn signal was demonstrated by the absence of a signal in the α-syn monomer-injected brains (**Fig. 2b** and **SVideo**. **4**). AOI987 was also tested for LSM but did not work with the iDISCO+ protocol (**SFig 5**). In addition, we demonstrated the use of TH and DAT antibodies for labeling dopaminergic neuron in the mouse brain following the adapted iDISCO+ protocol (**SFig 6**).

Next, we imaged α-syn aggregates at a resolution of approximately 1 µm within the intact cleared whole brain at 20× magnification to validate the accuracy of the pS129+ signal for α-syn aggregates in the striatum and SNc. We observed characteristic somatic and neuritic morphology of the α-syn aggregates in the LSM images (**Fig 2c, d**), similar to that typically observed with conventional immunohistochemical staining of 30-µm-thick brain sections (**SFig 2**). The specificity of signal detection by the automated analysis pipeline using the optimized iDISCO+ protocol was confirmed by analyzing the segmentation map with raw images acquired at 2× magnification (2.75×2.75 µm resolution) in the SNc and striatum (**Fig 2c–f**). A comparative analysis of representative somatic and neuritic aggregates revealed that imaging at 20× provided a sharper signal with a smaller full width at half maximum (**Fig 2c–f**). We used support vector machine-based detection and mapped the local density of α-syn aggregates throughout the entire brain. We revealed prominent labeling in the SNc, striatum, NAc, CEA, PAG and PPN (**Fig 2g, h**), corroborating the classical grading by a neuropathologist based on immunohistochemistry using free-floating sections.

### Automated high-accuracy whole-brain LSM-MRI image coregistration and tissue shrinkage assessment

Next, we developed an automated pipeline for the cross-scale integration of macroscopic MRI and microscopic LSM data (**SFig. 7**). The implemented pipeline employs a parallelized, multistage registration framework based on the ANTs (Advanced Normalization Tools) library to align the LSM and MRI data within a unified 20-µm coordinate space. We introduce an MRI-guided framework that integrates artifact masking and bidirectional transforms to quantify region-level clearing distortions. Leveraging MRI as a structural reference, the framework achieves robust registration while explicitly characterizing clearing-induced deformations. The framework achieved superior global alignment, with a mean MRI-LSM landmark error of 41.9±22.4 µm across seven landmarks in three iDISCO+-processed mice^28,29^ (**Fig 3**, **Table 1**). It maintained high precision in distortion-prone structures, including the ventricular system (lateral ventricle: 26.7±4.7 µm) and fiber-rich white matter tracts (olfactory limb: 23.3±4.7 µm). Visual inspection across multimodal anatomical alignment revealed consistent anatomical continuity. Structural accuracy and well-preserved topology were evidenced by low Jacobian determinant variability (6.60±0.18%), low mean squared error (0.0481±0.0073), and high local correlation (88.36±0.57%).

**Table 1.**
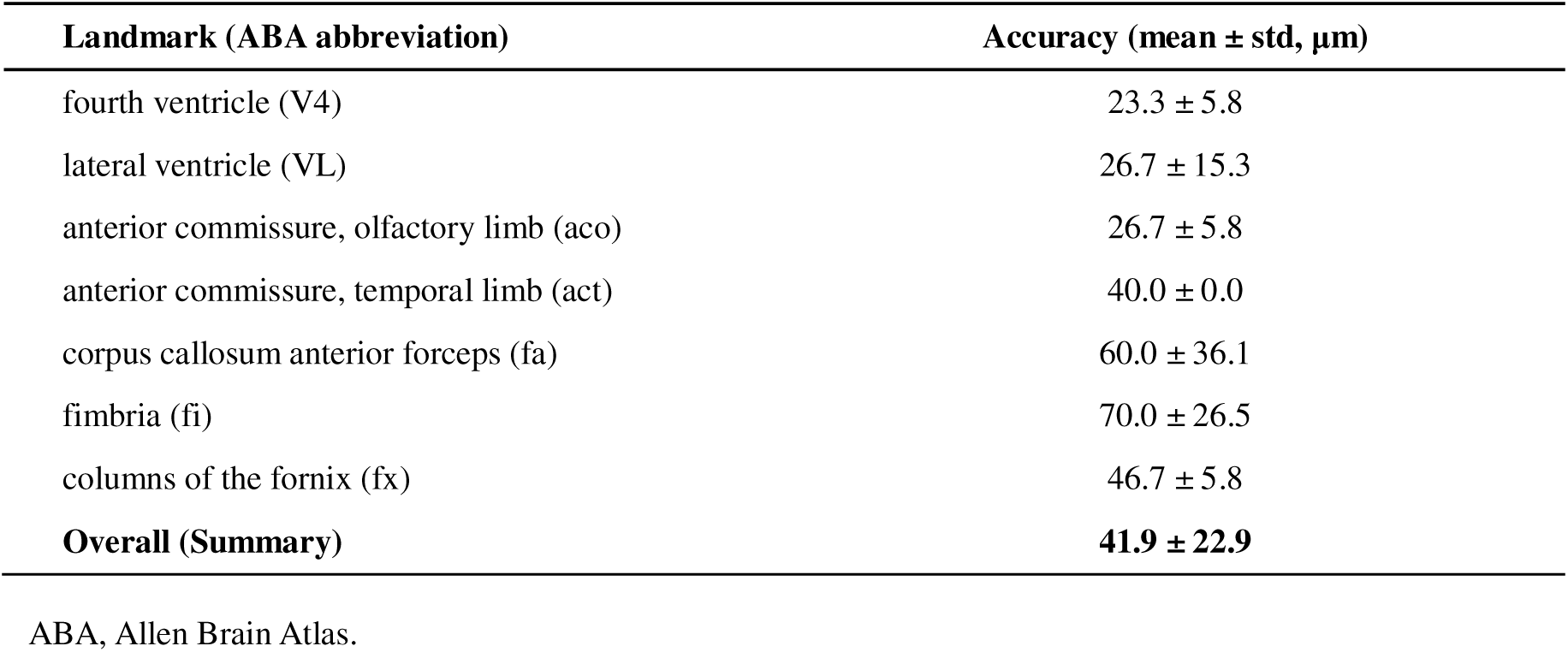
Landmark-based evaluation of co-registration accuracy.

| **Landmark (ABA abbreviation)** | **Accuracy (mean ± std, $\mu\text{m}$)** |
| --- | --- |
| fourth ventricle (V4) | 23.3 ± 5.8 |
| lateral ventricle (VL) | 26.7 ± 15.3 |
| anterior commissure, olfactory limb (aco) | 26.7 ± 5.8 |
| anterior commissure, temporal limb (act) | 40.0 ± 0.0 |
| corpus callosum anterior forceps (fa) | 60.0 ± 36.1 |
| fimbria (fi) | 70.0 ± 26.5 |
| columns of the fornix (fx) | 46.7 ± 5.8 |
| **Overall (Summary)** | **41.9 ± 22.9** |

ABA, Allen Brain Atlas.

Although the clearing protocol is known to induce tissue shrinkage (**SFig. 8**), the extent to which each brain region shrinks has not been evaluated. Here, we assessed global deformation and anisotropic scaling following tissue clearing. Quantitative evaluation of global tissue shrinkage was performed across the three principal macroscopic axes (X, Y, and Z). Subsequent deformation analysis indicated 9.32% global tissue shrinkage, with K-means clustering identifying stable, uniformly shrunk, and highly distorted regions. High anisotropy and distortion were concentrated in fiber-rich tracts and deep cortical layers, reflecting the heterogeneity of myelin and tissue density. This provides a standardized and quantitative basis for assessing morphological alterations and supports subsequent systematic, atlas-informed multimodal analysis of cleared brain samples.

### Whole-brain mapping revealed that **α**-syn PFF SNc injection led to the spread of **α**-syn pathology and dopaminergic degeneration

To assess the impact of α-syn PFF injection on the dopaminergic nigrostriatal pathway, we visualized p-α-syn deposits and TH-positive neuron across the whole brain using the optimized iDISCO+ protocol in PFF- and monomer-injected mice at 12 and 20 wpi (**Fig. 4a, b)**. We used the molecular atlas for the subregional analysis of the striatum (**Fig. 4a, SFig. 9**). pS129 levels were greater in different brain regions of PFF mice (at 12 and 20 wpi) than in those of the monomer group in the ipsilateral hemisphere (SNc, striatum, bed nuclei of the stria terminalis (BST), CEA, and VTA), except for no difference in the medial-posterior SNc or striatum (11) subregion at 20 wpi (**Fig 4, 5a–c, h**). An increase in pS129 levels in the contralateral hemisphere was observed only at 20 wpi in PFF mice compared with those in the SNc (especially the medial), striatum (06 and 11), and anterior VTA and BST but at both 12 and 20 wpi in the medial CEA (**Fig 4, 5a–d**). pS129 levels in the ipsilateral hemisphere remained stable at 12 wpi and 20 wpi in most striatal subregions, such as the BST, CEA and VTA (**Fig 4, 5a–c**), with a marked decrease in the ipsilateral SNc and an increase in the contralateral anterior VTA. These findings indicate that pathological α-syn spreads across the brain through the nigrostriatal pathway (**Fig 4, 5a–d**). Voxelwise spatiotemporal analysis also confirmed widespread increases in pS129 fluorescence intensity in the ipsilateral SNc, striatum, BST, and CEA in PFF-injected mice at 12 and 20 wpi compared with those in the monomer-injected group and differences in SNc between the 12 and 20 wpi PFF groups (**Fig. 8b, SFigs. 10**, **11**).

**Figure 4.**
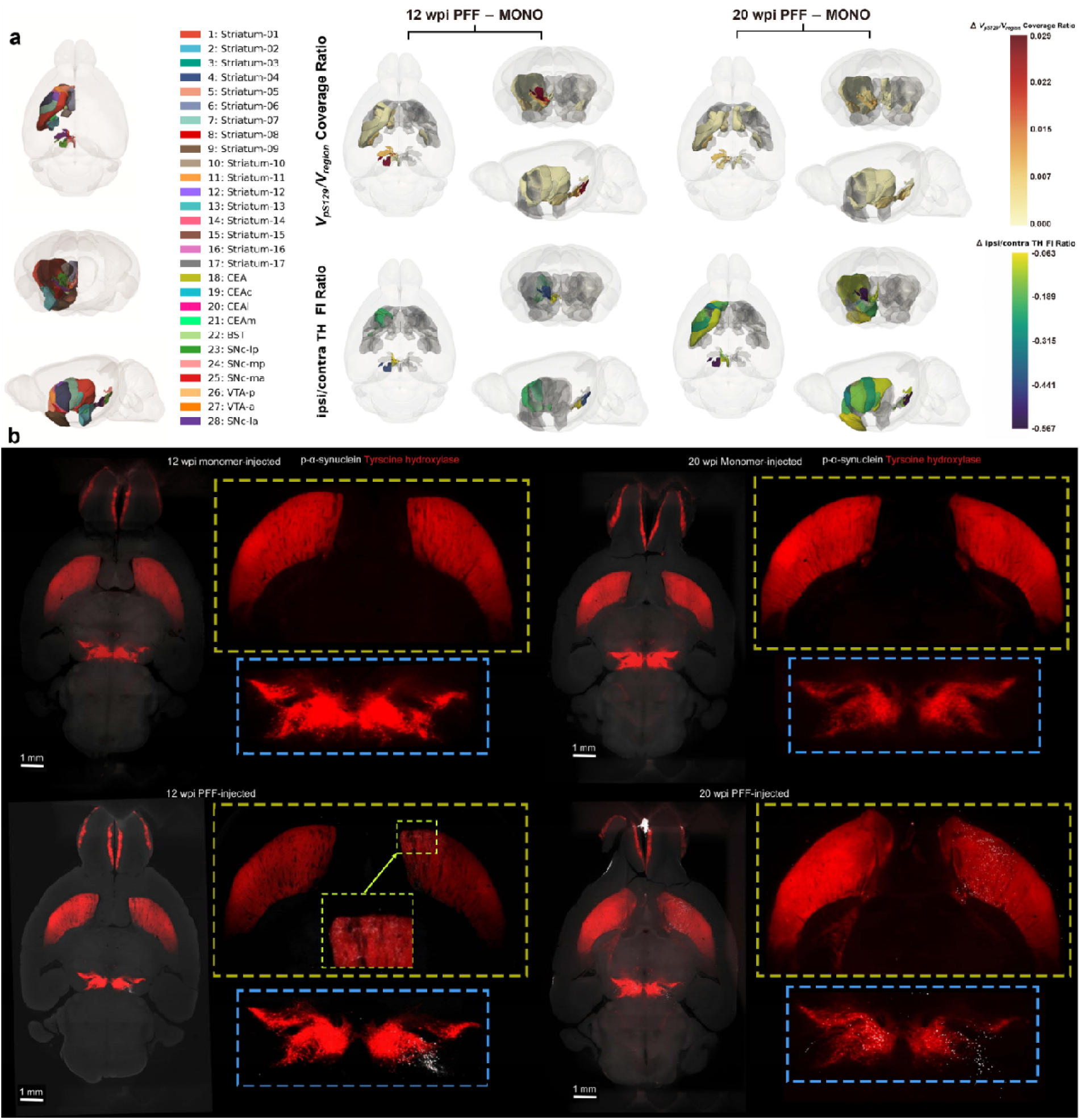
Neuroanatomical mapping and quantitative 3D visualization of α-synuclein pathology and dopaminergic integrity. **(a)** 3D neuroanatomical atlas of ROIs and differential rendering of systemic α-synuclein propagation and dopaminergic integrity. Smoothed surface mesh rendering depicting the 3D spatial locations of distinct ROIs included in the brain-wide quantitative analysis. ROIs were derived from three sources: the Allen Brain Atlas (ABA), a high-resolution Molecular Atlas for striatal sub-regions, and a dynamic consensus atlas subdividing the SNc and VTA based on endogenous TH distribution. A transparent whole-brain mesh is included for anatomical reference. Volumetric 3D visualization mapping the statistically significant differences in α -synuclein pathology load and dopaminergic marker intensity between PFF-injected and monomer-injected mice across different timepoints (12 and 20 wpi). Brain regions exhibiting significant differences (non-parametric Mann-Whitney U test, P < 0.05) are rendered and color-coded based on the median value difference (PFF minus Monomer). Non-significant brain regions are rendered in transparent silver to show comprehensive spatial context within a transparent whole-brain shell. **(b)** High-resolution MIP visualization of the nigrostriatal pathway. High-resolution Maximum Intensity Projection (MIP) of the spatial distribution of phosphorylated α-synuclein (pS129) and Tyrosine Hydroxylase (TH+, indicating dopaminergic neurons) along the nigrostriatal pathway. An inset displays a magnified view of the striatum region from the 12 wpi PFF-injected brain, where the brightness has been uniformly increased by 20% to better visualize the relatively sparse pS129 pathological distribution. Scale bar = 1 mm. ABA: Allen Brain Atlas, MIP: Maximum Intensity Projection, PFF: Preformed fibrils, pS129: Phosphorylated α-synuclein at Serine 129, ROI: Region of interest, SNc: Substantia nigra pars compacta, TH: Tyrosine hydroxylase, TH+: Tyrosine hydroxylase positive, VTA: Ventral tegmental area, wpi: Weeks post-injection.

**Figure 5.**
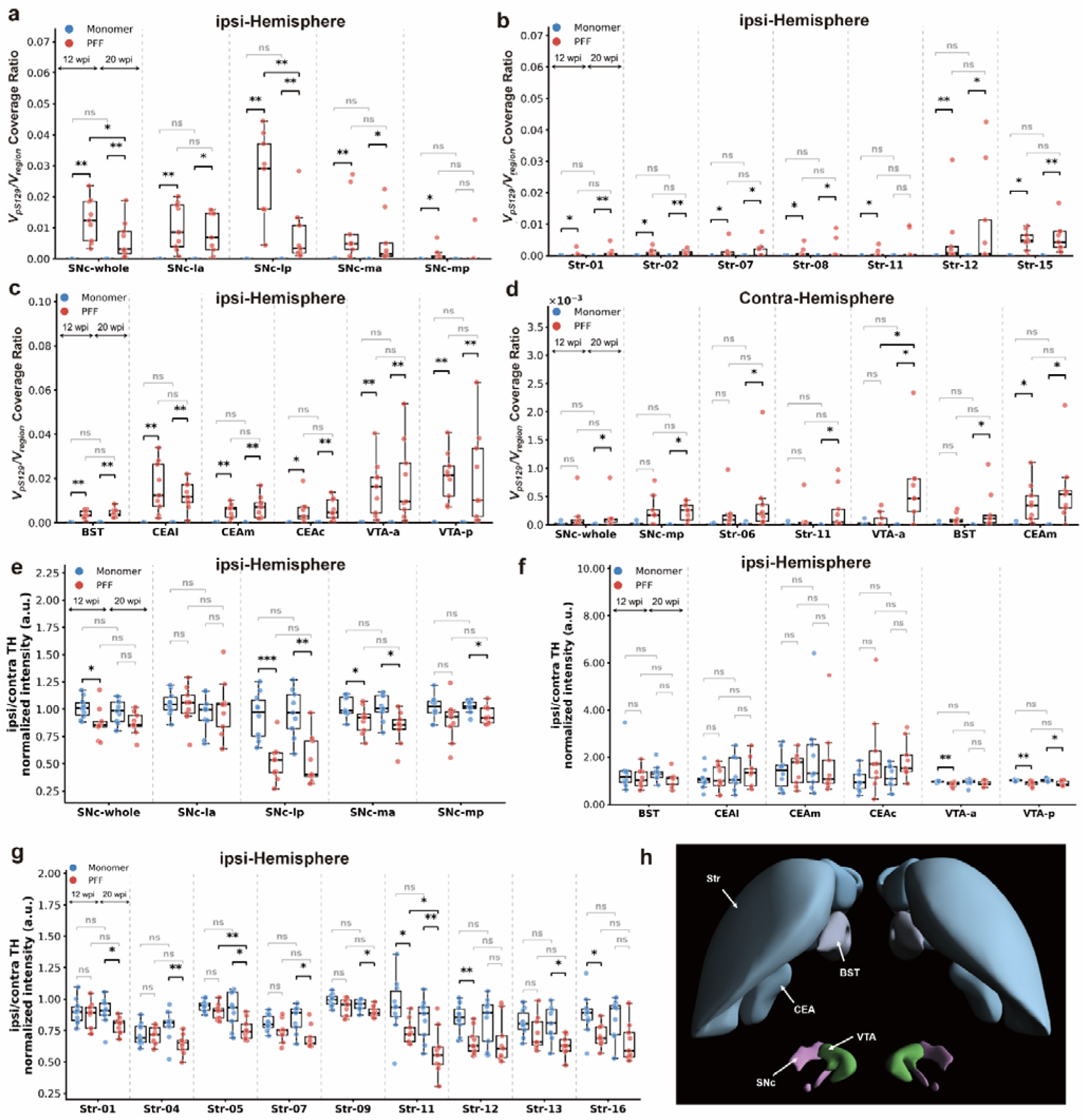
Regional comparisons of pS129 aggregate volumetric coverage and TH fluorescence intensity across monomer-injected controls and PFF-injected mice at 12- and 20- wpi. **(a-c)** Assessment of ipsilateral (left hemisphere) pS129 coverage ratio (V_pS129_ /V _region_) across finely parcellated anatomical sub-regions, including (a) the substantia nigra pars compacta (SNc), (b) distinct striatal sub-regions, and (c) extended interconnected nodes (amygdaloid complex and ventral tegmental area). **(d)** Quantification of contralateral (right hemisphere) pS129 coverage ratio in selected target regions. **(e-g)** Evaluation of dopaminergic integrity via the ipsilateral-to-contralateral (L/R) TH intensity ratio within the corresponding sub-regions, including (e) the SNc, (f) extended interconnected nodes (amygdaloid complex and ventral tegmental area), and (g) distinct striatal sub-regions. Statistical significance between designated groups was determined using the two-sided non-parametric Mann-Whitney U test. *P < 0.05, **P < 0.01, ***P < 0.001, ns = not significant. **(h)** Three-dimensional anatomical structures and spatial distributions of the selected ROIs. Renderings were generated using the Allen Brain 3D Explorer (https://connectivity.brain-map.org/3d-viewer). BST: Bed nuclei of the stria terminalis, CEAc: Central amygdalar nucleus, capsular part, CEAl: Central amygdalar nucleus, lateral part, CEAm: Central amygdalar nucleus, medial part, PFF: Preformed fibrils pS129: Phosphorylated α-synuclein at Serine 129, ROI: Region of interest, SNc: Substantia nigra pars compacta, SNc-la: SNc, lateral anterior, SNc-lp: SNc, lateral posterior, SNc-ma: SNc, medial anterior, SNc-mp: SNc, medial posterior, Str: Striatum (numbered 01 to 16 based on molecular atlas sub-parcellation), TH: Tyrosine hydroxylase, VTA-a: Ventral tegmental area, anterior, VTA-p: Ventral tegmental area, posterior, wpi: Weeks post-injection.

3D spatial renderings of the bi-hemispheric pS129 covariance networks and hierarchical chord diagrams illustrated significant trans-hemispheric and intra-hemispheric links (**Fig. 7d, e**). Covariance links were observed in the ipsilateral macroscopic covariance networks of pS129 pathology at 12 and 20 wpi in PFF-injected mice, especially for the striatum, SNc, CEA and BST, with the connecting edges denoting significant pairwise Pearson correlations of regional pS129 coverage (**Fig. 7d**).

Compared with those in the ipsilateral hemisphere (medial-anterior, medial-posterior and lateral-posterior SNc, Striatum 11, and posterior VTA), the intensity levels in the ipsilateral hemisphere (at 12 and 20 wpi) in the ipsi/contra TH-normalized group were reduced (**Fig 4, 5e–g**). In addition, the normalized ipsi/contra TH intensity in the ipsilateral hemisphere was lower in the striatal subregion (5 and 11) in 20 wpi PFF mice than in 12wpi PFF mice (**Fig 4, 5g**). The reduction in the normalized ipsi/contra TH intensity in the striatum and SNc was further confirmed by manual analysis of TH-positive neuron counts in the 12 wpi group (**SFig 12**). Voxelwise spatiotemporal analysis also confirmed a widespread reduction in TH fluorescence intensity in the ipsilateral as well as in the contralateral SNc and striatum of PFF-injected mice at 12 and 20 wpi compared with that in the monomer-injected group and the differences between the 12 and 20 wpi PFF groups (**Fig. 8b, SFigs. 10, 11**). However, a decrease in the normalized ipsi/contra TH intensity was not observed in the contralateral hemisphere at 12 or 20 wpi in the PFF groups compared with the monomer groups. Further regional cross-modal correlation analyses revealed correlations between the pS129 coverage ratio (Vp5129/Vregion) and the normalized ipsi/contra TH intensity within the lateral-posterior SNc and posterior VTA across all groups (**Fig 7a**). The temporal evolution matrix illustrated the normalized median differences between α-syn PFF-injected mice and monomer-injected mice at 12 and 20 wpi (**Fig 8a**). This demonstrated the direction and magnitude of the alterations in pS129 and TH across whole-brain regions due to α-syn PFF injection in the SNc.

### α-syn PFF SNc injection leads to microstructural and iron-related alterations in the brains of PFF-injected mice

Next, we assessed *the* neuroanatomical, microstructural and iron-related changes in the mouse models *by using ex vivo* T1w, DTI, and SWI MRI (9.4T). DTI quantifies water diffusion to assess microstructural integrity and can reveal impaired white matter integrity in dopaminergic nuclei and motor pathways, indicating widespread structural disruptions. Voxelwise group comparisons of T1w DBM as well as DTI DBM measures were performed using permutation-based *t* tests with TFCE. Comparisons included α-syn monomer- and PFF-injected mice at 12 and 20 wpi, as well as longitudinal within-group comparisons. Significant effects were observed for both expansion and atrophy in the monomer-injected mice at 12 vs. 20 wpi and for regional atrophy in the PFF-injected mice at 12 vs. 20 wpi, whereas no other comparisons reached statistical significance after correction (TFCE-corrected *p* < 0.05) (**STable 2, SFig 13**).

Representative coronal ex vivo AD, MD, RD, and FA maps of monomer- or PFF-injected mice at 12 and 20 wpi, acquired at 100 μm isotropic resolution via DTI MRI. Compared with those in the control mice, the RDs in the striatal subregions (05 and 10 defined by the molecular atlas^30^) and posterior VTA in the PFF-injected mice at 20 wpi were decreased (**Fig. 6d)**. Increased AD (ipsilateral brachium of the superior colliculus and contralateral dorsal hypothalamus) and MD (contralateral dorsal hypothalamus) and decreased AD and MD in the striatal subregion (17) were observed in PFF-injected mice at 20 wpi compared with those in the 12wpi group (**Fig. 6a, b**; **STable 3**).

**Figure 6.**
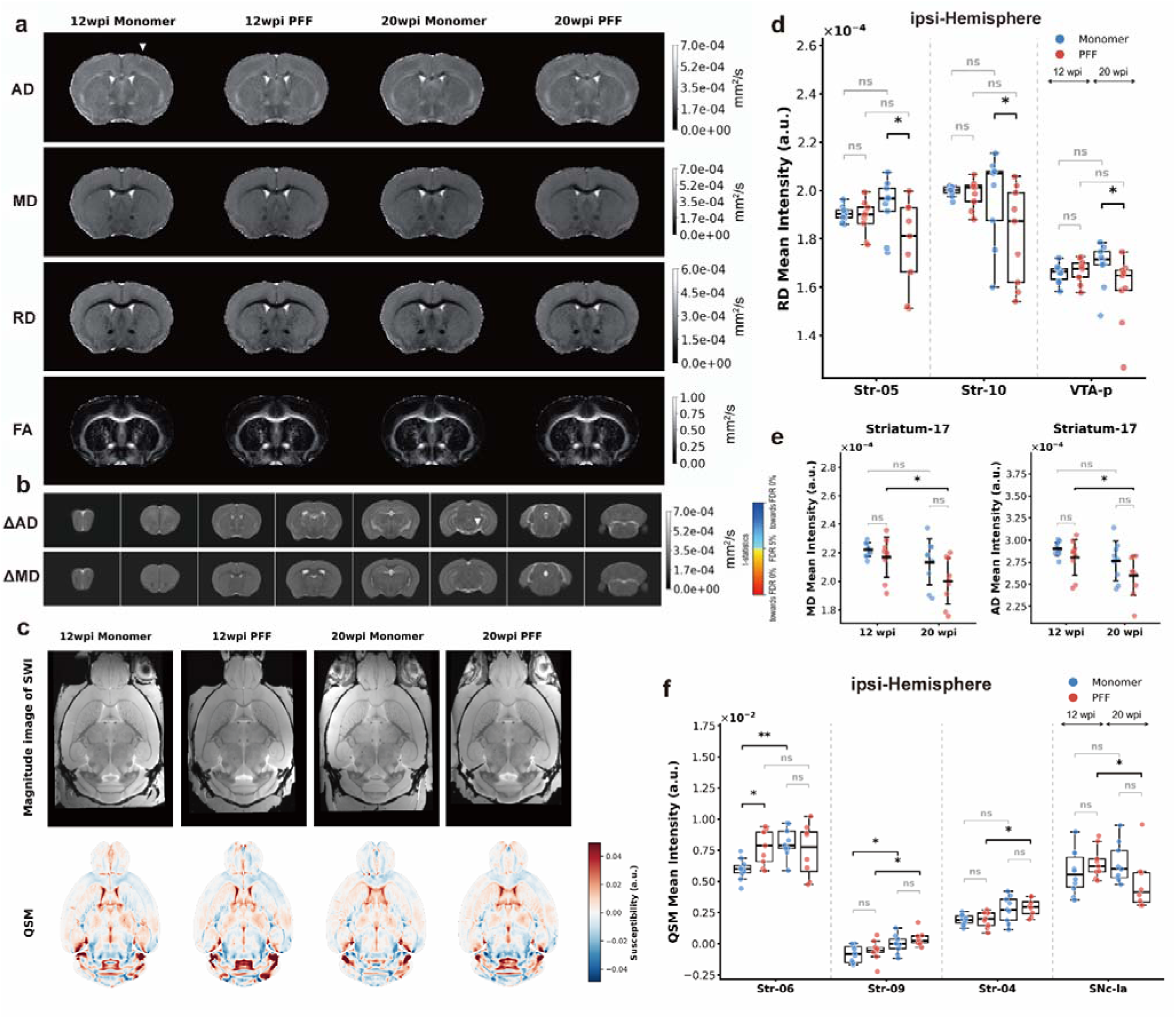
Comprehensive multi-parametric MRI evaluation reveals microstructural and compositional alterations in the brains of PFF-injected mice. **(a)** Representative coronal ex vivo AD, MD, RD, and FA maps of monomer- or PFF-injected mice at 12 and 20 wpi, acquired at 100 μm isotropic resolution via DTI MRI. The white triangle indicates the injection side. **(b)** Voxel-wise differences in AD and MD between PFF-injected mice at 12 and 20 wpi. Voxel-wise analyses of AD and MD are shown as t-statistic maps overlaid on the anatomical template for the comparison 12 wpi PFF vs. 20 wpi PFF. Red/yellow colors indicate regional increases in the corresponding diffusion metric. Significant increases in AD were observed in the ipsilateral brachium of the superior colliculus and contralateral dorsal hypothalamus, while MD increases were detected in the contralateral dorsal hypothalamus. The color bar represents t-statistics, thresholded using TFCE-corrected permutation testing (p < 0.05). The white triangle represents the injected side. **(c)** Representative horizontal section of SWI magnitude images (top row) and their corresponding reconstructed QSM maps (bottom row) across the four experimental groups. **(d)** Significant reductions in RD mean intensity within striatal and posterior VTA regions in the PFF cohort at 20 wpi relative to monomer controls. **(e)** Regional variations in MD and AD mean intensity across specific striatal sub-regions in the PFF-injected cohort defined by the molecular atlas. **(f)** Alterations in QSM mean intensity evaluated across targeted striatum and SNc sub-regions, reflecting variations in tissue magnetic susceptibility. Statistical significance was calculated using the two-sided non-parametric Mann-Whitney U test. *P < 0.05, **P < 0.01, ***P < 0.001, ns = not significant. α-syn: α-synuclein, AD: Axial diffusivity, DTI: Diffusion tensor imaging, FA: Fractional anisotropy, MD: Mean diffusivity, PFF: Preformed fibrils, QSM: Quantitative susceptibility mapping, RD: Radial diffusivity, ROI: Region of interest, SNc-la: Substantia nigra pars compacta, lateral anterior, Str: Striatum, SWI: Susceptibility-weighted imaging, TFCE: Threshold-free cluster enhancement, VTA-p: Ventral tegmental area, posterior, wpi: Weeks post-injection.

Additionally, the mean intensity of the QSM, reflecting tissue magnetic susceptibility (Mann-Whitney U test), was greater in the striatal subregion (06) in 12 wpi PFF-injected mice than in monomer mice. The mean intensity of the QSM was lower in the lateral anterior SNc of 20 wpi PFF mice than in that of 12 wpi PFF mice but was greater in the striatal subregion (04 and 09) (**Fig. 6c, f**). Further regional cross-modal correlation analyses revealed correlations between the pS129 coverage ratio (Vp5129/Vregion) and the mean QSM intensity in the SNc and medial CEA in 12 wpi PFF mice, and the mean QSM intensity and ipsi/contra TH normalized intensity in the striatal subregions (13 and 15) when all PFF-injected mice were grouped (**Fig. 7b, c**). The temporal evolution matrix of changes in QSM and DTI scalars across whole-brain regions further demonstrated the magnitude of the alterations due to α-syn PFF injection in the SNc.

**Figure 7.**
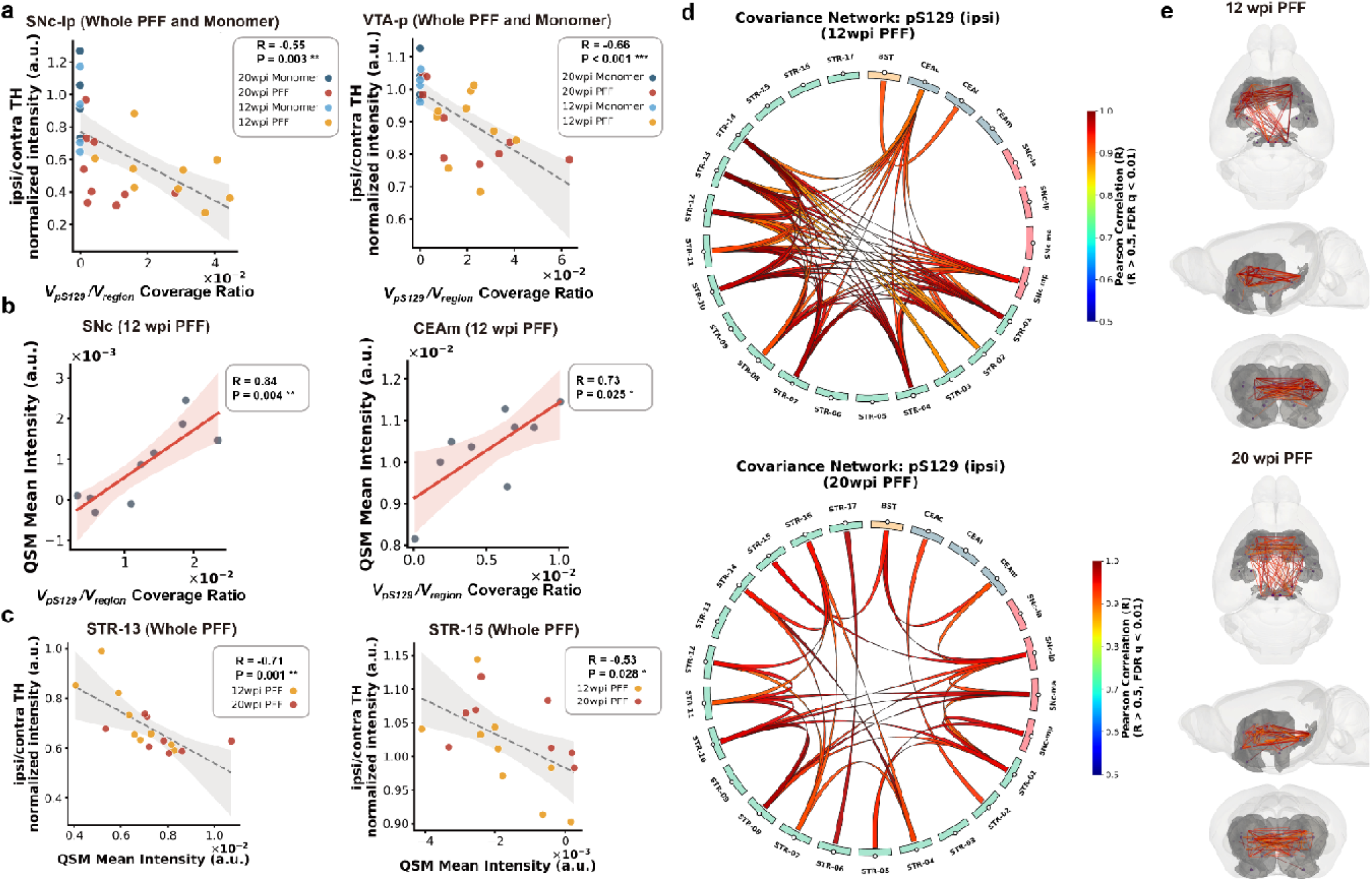
Cross-modal covariance analysis and correlation-based association network mapping of α-synuclein propagation and TH^+^ neurodegeneration. **(a–c)** Regional cross-modal correlation analyses evaluating the linear associations between distinct pathological and microstructural metrics. Scatter plots demonstrate the correlations between **(a)** the pS129 coverage ratio () and the ipsilateral/contralateral (ipsi/contra) TH normalized intensity within the SNc-lp and VTA-p across all cohorts, **(b)** the pS129 coverage ratio and QSM mean intensity in the SNc and CEAm for the 12 wpi PFF cohort, and **(c)** QSM mean intensity and ipsi/contra TH normalized intensity in striatal sub-regions (STR-13 and STR-15) for the combined PFF cohort. Statistical significance was assessed using the Pearson correlation coefficient (R) and shaded regions denote the 95% confidence intervals. **(d)** Hierarchical chord diagrams illustrating the ipsilateral macroscopic covariance networks of pS129 pathology at 12 and 20 wpi in PFF-injected mice. The outer ring nodes represent fine-grained anatomical regions of interest (ROIs), color-coded by macro-structural identity (i.e., green for striatal subdivisions, red for SNc subdivisions, blue for CEA, and orange for BST). Connecting edges denote significant pairwise Pearson correlations of regional pS129 coverage. To ensure robust association mapping, network edges were strictly thresholded to retain only strong positive correlations (R > 0.5) that survived Benjamini-Hochberg false discovery rate (FDR) multiple comparison correction (q < 0.01). **(e)** 3D spatial renderings of the bi-hemispheric pS129 covariance networks. Significant trans-hemispheric and intra-hemispheric covariance links (filtered using the identical rigorous thresholds: Pearson R > 0.5, FDR q < 0.01) are projected directly into the 3D anatomical reference space. BST: Bed nuclei of the stria terminalis, CEA: Central amygdalar nucleus (c: capsular, l: lateral, m: medial), FDR: False discovery rate, ipsi/contra: Ipsilateral to contralateral ratio, PFF: Preformed fibrils, pS129: Phosphorylated α-synuclein at Serine 129, QSM: Quantitative susceptibility mapping, ROI: Region of interest, SNc: Substantia nigra pars compacta (la: lateral anterior, lp: lateral posterior, ma: medial anterior, mp: medial posterior), STR: Striatum (numbered subdivisions based on the molecular atlas), TH: Tyrosine hydroxylase, VTA: Ventral tegmental area (a: anterior, p: posterior), wpi: Weeks post-injection.

**Figure 8.**
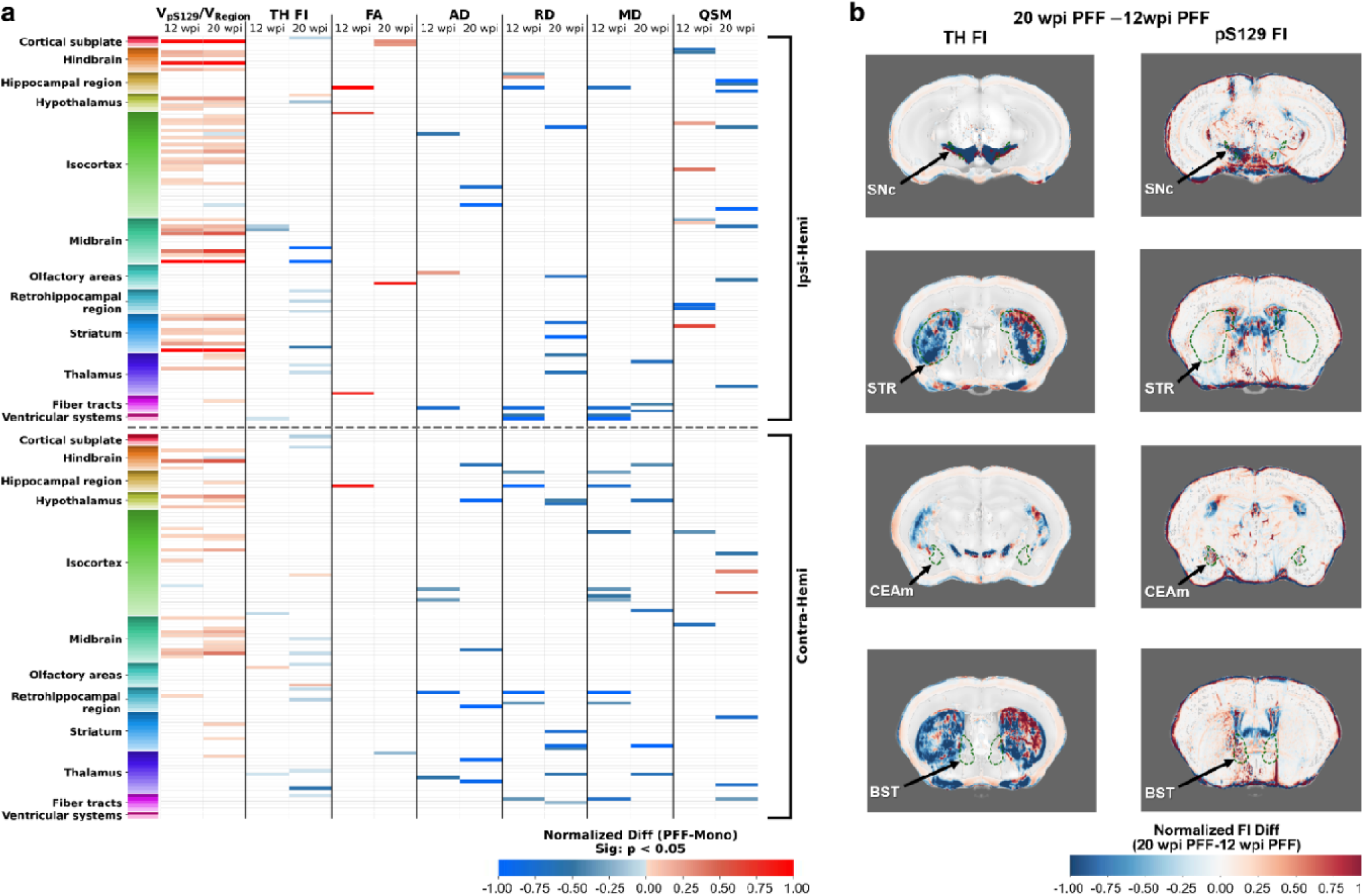
Spatiotemporal profiling of whole-brain multimodal metrics and voxel-wise TH alterations in the α-syn PFF mouse model. **(a)** Temporal evolution matrix of multimodal neuropathological changes across whole-brain regions at 12 and 20 wpi. The matrix displays the normalized median differences between α-syn PFF-injected mice and monomer-injected controls. Rows represent specific brain ROIs, hierarchically grouped by major anatomical categories (indicated by the left color-coded annotation bar) and physically segregated into ipsilateral (top) and contralateral (bottom) hemispheres. Columns indicate the evaluated multimodal metrics, including histological readouts (pS129 coverage, TH mean intensity) and neuroimaging parameters derived from DTI (FA, AD, RD, MD) and QSM. The color gradient denotes the direction and magnitude of the alterations. To strictly highlight robust disease-associated signatures, the heatmap is masked by statistical significance; white (blank) cells represent non-significant regional alterations (two-sided Mann-Whitney U test, p < 0.05). **(b)** Voxel-wise difference maps illustrating dopaminergic alterations assessed by TH expression for longitudinal comparisons within the PFF group (20 wpi vs. 12 wpi). Visualizations are generated using a MIP across a 10-slice window and overlaid onto an averaged tissue autofluorescence template. Warm colors denote increased pS129 burden. Green dashed contours delineate predefined anatomical boundaries of critical ROIs derived from the Allen brain atlas. α-syn: α-synuclein, AD: axial diffusivity, DTI: diffusion tensor imaging, FA: fractional anisotropy, MD: mean diffusivity, MIP: Maximum intensity projection, PFF: preformed fibrils, pS129: phosphorylated serine 129, QSM: quantitative susceptibility mapping, RD: radial diffusivity, ROI: region of interest, TH: tyrosine hydroxylase, wpi: weeks post-injection.

## Discussion

Here we developed an open-source, automated analysis for registering high-field MRI and LSM with <40 µm alignment error, multimodal multiscale brain-wide regional and voxel based analysis, also in the subregions of striatum. We adapted the IDISCO+ protocol to map the spatiotemporal evolution of α-syn pathology, dopaminergic degeneration by LSM at single-cell resolution, achieving uniform whole-brain labeling, including the densely innervated striatum. The specificity was confirmed by the absence of signal in monomer-injected controls and by high-magnification imaging that resolved characteristic somatic and neuritic aggregate morphology. Using this pipeline, we demonstrated that unilateral SNc PFF injection drives the spread of α-syn pathology along the nigrostriatal pathway to ipsilateral striatum, BST, CEA, and contralateral SNc. pS129 pathology in the ipsilateral SNc declines at 20 wpi compared to12 wpi, along with progressive loss of TH-positive neuron; In addition, α-syn pathology lead to region-specific microstructural and iron-related alterations that correlate with loss of TH-positive neuron and loss of TH-positive neuron.

Accurate cross-modal registration between LSM and MRI has historically been challenging due to differences in contrast mechanisms, resolution, and tissue clearing-induced nonlinear deformations. Previous studies used manual alignment, affine-only transformations, or did not systematically quantify clearing artifacts have not yet allowed automated registration and quantification^31–36^. Our pipeline addresses these gaps by implementing a parallelized, multistage diffeomorphic registration framework (ANTs) with bidirectional transforms, explicit artifact masking, and automated landmark-based validation. We achieved a mean landmark error of 41.9±22.4 µm and, quantified region-level clearing-induced distortion across the whole brain. We identified stable, uniformly shrunk, and highly distorted regions due to the adapted IDISCO+ protocol. This framework is openly available and can be readily used for analysis of other mouse brain LSM (with different clearing protocols), and MRI imaging data. This will facilitate the integrating microscopic pathology with MRI readouts of different resolution. While the spatial distribution of TH and pS129 closely matched the anatomical boundaries of the ROIs related to dopaminergic degeneration and α-Syn spread theories in the PFF mouse model, statistically significant voxels were restricted to localized clusters within the SNc and striatum. This limited voxel-level significance is likely a consequence of LSM downsampling and spatial averaging during image registration, which enhances morphological stability at the expense of localized statistical power. This highlight the importance to integrate voxelwise analysis with region- and subregion-level quantifications to capture the full pathological spectrum.

PFF-injected rodent models have been used to understand α-syn spread and its associated effects. In mice, the intracerebral, intraperitoneal, intramuscular, intravenous, oral administration, and kidney, gut-induced mouse or human α-syn PFFs results in widespread α-syn pathology in the brain in both wildtype and genetically modified mice ^10,13,37–41^ and in nonhuman primates^17,42^. These injections induce the misfolding and aggregation of endogenous α-syn, leading to its propagation along anatomically connected pathways, with the distribution pattern dependent on multiple factors including the site of initial seeding, and α-syn PFF injected. The distribution of α-syn pathology in mouse brain in the current study is in line with the earlier studies by Geibl et al ^10^ using the same mouse strain and α-syn PFF preparation protocol. Dadgar-Kiani *et al*. reported the α-syn spreads mostly in the temporal cortex, hypothalamus, posterior striatum and posterior midbrain at two and four months after bilateral mouse PFFs injection in the SNc ^25^. Masuda-Suzukake et al reported α-syn spreading to the ipsilateral striatum, hippocampus, amygdala, temporal cortex, and hypothalamus of mouse brain at 3 months after human PFF injection^37^.

One of the main finding is that dense ipsilateral SNc pS129 pathology at 12 wpi markedly decreased by 20 wpi, while TH+ neuron loss progressed during the same interval. This inverse relationship suggests that pS129 epitope loss may occur in severely degenerating neurons, either through aggregate compaction, proteolytic degradation, or neuronal death itself. This observation has critical implications for interpreting ongoing α-syn PET imaging studies in PD patients ^17,43,44^. The absence of detectable pathology at late stages does not necessarily indicate recovery but may instead reflect end-stage neurodegeneration. Our temporal evolution matrix and voxelwise analyses further showed that pS129 levels remained stable or increased in projection targets (striatum, BST, CEA) and the contralateral SNc, supporting a prion-like propagation model where pathology spreads antegradely and retrogradely along connected circuits. The distribution of pS129 pathology in our model (ipsilateral SNc -striatum (subregions)- BST/CEA - contralateral SNc) closely recapitulates the topographic pattern reported by Geibl et al.^10^ using the same PFF preparation, and extends it to a whole-brain subregional quantitative analysis.

Unlike aggressive neurotoxin-based models (MPTP, 6-OHDA), which cause rapid and nearly complete dopaminergic loss (e.g., 82% loss at 7 days post-MPTP), the PFF model induces a slower progression with partial denervation, approximately 30% TH loss by 20 wpi in our study. Moreover, we demonstrated quantitatively that the subregions are affected differently.This reduction in striatal TH is consistent with earlier findings of Geibl et al ^10^. PD patients typically exhibit approximately 50% reduction in striatal TH at diagnosis^6^. Thus the milder, progressive phenotype of the PFF model makes it more suitable for testing disease-modifying therapies and for studying the α-syn induced degeneration. Ina addition, we found decreased SNc pathology with ongoing TH loss challenges the simplistic view that α-syn burden directly correlates with neuronal survival—a nuance increasingly recognized in human PD, where Lewy body density does not always match neurodegeneration severity. Recent studies have shown that synaptic enrichment of pSer129 α-syn correlates with dopaminergic denervation in early-stage PD, suggesting that pathology at synaptic terminals may be more directly pathogenic than somatic aggregates.

Additionally, we found elevated striatal susceptibility at 12 wpi in PFF mice and dynamic changes between 12 and 20 wpi (decreased lateral anterior SNc, increased striatal subregions 04 and 09). Cross-modal correlation analyses revealed that pS129 coverage correlated positively with QSM in SNc and medial CEA at 12 wpi, while TH loss correlated with QSM in striatal subregions when all PFF mice were grouped. Here we found only minor volumetric changes using T1w in PFF and monomer injected mice. PFF-injected mice at 20 wpi showed decreased RD in striatal subregions and posterior VTA compared to monomer group, which suggests myelin compaction or reduced extra-axonal space. Increased AD/MD in superior colliculus/hypothalamus and decreased AD/MD in striatal subregion in 20wpi vs 12wpi PFF mice indicates region-specific microstructural reorganization, and a combination of axonal injury, neuroinflammation, and compensatory plasticity. Previous studies in rodent PD models has detected microstructural changes following neurotoxin or PFF injection ^4619^, such as decreased FA and increased MD in MPTP or rotenone SN injected model^47^ and intramuscular α-syn PFF injected model^48^. The MitoPark model also shows reduced FA in the SN and corpus callosum^61^. Our study further providing whole-brain DTI at a higher resolution (100 µm isotropic) and by directly correlating DTI scalars with pS129 pathology and TH loss from the same animals enabled by multimodal registration. In human PD, studies showed that nigral iron accumulation and a “V-shaped” striatal iron trajectory^45^ associated with pathological progression. Dopaminergic striatal dysfunction and cell loss in the substantia nigra pars compacta are interrelated with increased nigral iron content ^49^. structural MRI using T1w morphometry reveals subtle macroscopic changes, including cortical thinning and basal ganglia, subcortical gray matter reductions, even in early disease^1514^, and large multicenter studies have shown that stage-specific alterations in the putamen, caudate, thalamus, and hippocampus correlate with motor and cognitive severity^16^. Microstructural changes has been observed in PD patients at different stages and was found associated with early motor dysfunction^49–52^.

This study has several limitations. 1) We used mouse α-syn PFFs in the study, mouse α-syn fibrils are known to be structurally and functionally distinct from human PFF ^53^. 2) whether α-syn spread antegrade or retrograde, and cell-to-cell transfer mechanism has not be assessed in the current study^54,55^. Third, ex vivo fixation and clearing alter tissue morphology and diffusivity, although all groups were processed identically, and we quantified clearing-induced distortion to enable correction 3) We used ex vivo MRI rather than in vivo scans in the current study. However ex vivo MRI eliminates motion artifacts from breathing or involuntary movements, enabling longer scan times and higher resolution and signal-to-noise ratio.

In conclusion, our open-source pipeline and optimized iDISCO+ protocol allow for integrating cellular pathology with MRI-based biomarkers in mouse models. By enabling precise, multiscale automated mapping, this study facilitate better understanding of the mechanism underlying α-syn spread leading to microstructural changes and neurodegeneration in synuclinopathy model. This platform can be applied to assess candidate disease-modifying therapies (e.g., anti-α-syn antibodies) in a quantitative, whole-brain manner, and for general registration between MRI and LSM data.

## Materials and methods

### Animals

All animal experiments involving α-syn PFF injections were performed according to the ethical standards of the institution at which the studies were conducted (Regierungspräsidium Giessen, Germany V54-19c2015h01GI 20/28 Nr. G32/2022). Research was conducted according to the ARRIVE 2.0 Guidelines. The mice were housed in groups of up to 5 animals per cage with food and water provided ad libitum on a 12-h light/dark cycle. All the experiments were performed in the light phase. A total of 46 C57BL/6 J mice (Charles River, Sulzfeld, Germany) were used; 5 PFF-injected mice, including 22 PFF- (4 for protocol optimization) and 19 monomer-injected mice, were used for whole-brain immunohistochemical analysis of pS129 pathology assessment and 41 for the LSM experiments. A similar distribution of male and female mice was used in all the experiments. All the animals were between two and three months old at the beginning of the experiments. Six additional male B6129SF2/J mice, 14–15 months old (F2 hybrid strain; The Jackson Laboratory, USA), were used for preliminary protocol optimization.

### Stereotaxic injections of purified **α**-syn PFFs and **α**-syn monomers

Mouse full-length α-syn PFFs were prepared at Johns Hopkins University School of Medicine, Baltimore, USA, as previously described ^10–12^ (details in the supplemental file). For the stereotactic delivery of α-syn PFFs and monomers, the mice were anesthetized with fully antagonizable anesthesia containing medetomidine (500 µg/kg body weight), midazolam (5 mg/kg body weight), and fentanyl (50 µg/kg body weight). A total volume of 1 µl of α-syn PFFs or monomers was divided and injected into two injection sites in the SNc (anteroposterior (AP): - 3.10; mediolateral (ML): -1.30; dorsoventral (DV): -4.45; and AP: -3.50; ML: -1.30; DV: -4.20), intended to cover the entire region (**SFig. 1**). All injections were performed unilaterally into the left hemisphere using a microinjector at a velocity of 125 nl/min. At the end of the surgery, the anesthesia was antagonized with atipamezole (2.5 mg/kg body weight) and flumazenil (0.5 mg/kg body weight).

### Tissue processing

The mice were anesthetized with a mixture of ketamine (50 mg/kg) and xylazine (4.5 mg/kg) and sacrificed by transcardial perfusion with ice-cold 0.1 M PBS (pH 7.4) and 4% paraformaldehyde in 0.1 M PBS. For the immunohistochemical analysis of α-syn propagation using 30 µm-thick coronal sections, the mice were decapitated, and the brains were quickly removed, followed by postfixation for 3 days in paraformaldehyde solution and 3 days in 30% sucrose solution in 0.1 M PBS. The brains were then frozen on dry ice and stored at -80°C until sectioning. On the day of sectioning, the brains were embedded in tissue freezing media (OCT Compound, Tissue Tek, USA) and cut into 30 μm-thick consecutive coronal sections using a cryostat microtome (CM3050 S, Leica, Germany)^56,57^. All sections spanning the complete rostrocaudal extent of the brain were kept in the correct order and stored at 4°C in cryoprotect solution (1:1:3 volume ratio of ethylene glycol:glycerol:0.1 M PBS) until further processing. For the MRI and brain clearing experiments, the mice were decapitated, and complete heads (brains in the skull) were fixed in paraformaldehyde solution for 5 days. Afterward, complete heads were stored intact in 0.1 M PBS + 0.05% NaN_3_ and transferred to the University of Zurich and stored in 0.1 M PBS + 0.05 NaN_3_ (pH 7.4) at 4°C for 2 to 4.5 months before MRI scanning, as recommended.^27^ The brains were not removed from the skull to preserve cortical and central brain structure.

### Ex vivo high-field MR image

We took advantage of the extended data acquisition times available in *ex vivo* DTI to achieve high spatial resolution, avoiding image corruption from motion or breathing, which are issues commonly encountered in in vivo DTI, particularly in deep structures such as the SNc. Mouse head samples were placed in a 5 ml Eppendorf tube filled with Perfluoropolyether YR-1800 (GALDENR HT270, Apollo Scientific). Data were acquired on a BioSpec 94/30 preclinical MRI scanner (Bruker BioSpin AG, Fällanden, Switzerland) with a cryogenic 2×2 radio frequency phased-array surface coil (overall coil size of 20×27 mm^2^). The coil system operated at 30-K for reception in combination with a circularly polarized 86 mm volume resonator for transmission ^58,59^. A structural T1-weighted (T1w) scan was obtained using a 3D fast low-angle shot (FLASH) gradient echo sequence, with an echo time (TE) of 35 ms and a repetition time (TR) of 97 ms. The field of view (FOV) was set to 16.8×11.6×7.8 mm, with a matrix dimension of 280×194×130, resulting in a nominal voxel resolution of 60×60×60 μm. The total acquisition times were approximately 1 hour and 30 minutes. DTI was performed with a 3D multishot echo-planar imaging (EPI) sequence (4 shots) with an FOV of 20.9×11.6×7.8 mm and a matrix dimension of 209×116×78, resulting in a nominal voxel resolution of 100×100×100 μm. The following imaging parameters were used: a TR of 1500 ms, a TE of 28 ms, and no averaging. A total of 5 volumes were acquired, including one with a b value of 0 (A0) and four with a b value of 4000 s/mm², distributed across 60 diffusion encoding directions. A global and MAPSHIM protocol with a field map (default settings) was used for shimming. The DTI acquisition time was approximately 8 hours 30 minutes ^60^. SWI data were collected using a three-dimensional gradient-echo FLASH sequence with a TR of 250 ms and a TE of 12 ms. Four signal averages were acquired. The FOV was 16.8×11.6×7.8 mm with a matrix size of 279×194 ×130, resulting in an isotropic voxel resolution of approximately 60 μm. Fat suppression and outer-volume saturation were applied to reduce unwanted signal contributions. A global and MAPSHIM protocol with a field map (default setting) was also used for shimming. The total SWI acquisition time was approximately 7 hours ^61^. Details on the MRI data processing and analysis are provided in the supplemental files.

### Whole-brain tissue clearing and fluorescence labeling

The brains were removed from the skulls, immunolabeled and cleared using the classical iDISCO+ protocol ^62^. Briefly, the tissues were first dehydrated through a series of methanol solutions and then bleached in methanol containing 5% hydrogen peroxide (H_2_O_2_). After rehydration, the brains were permeabilized and blocked with 0.2% gelatin instead of NDS as previously described. The tissues were then incubated with a rabbit anti-pS129 α-syn primary antibody (Abcam, ab51253, 1:1000) for 1 week at 37°C. After multiple washes, the brains were incubated for 1 week at 37°C with anti-rabbit Cy3 secondary antibody (Jackson, 111-165-144, 1:1000) to reveal pS129 α-syn, together with conjugated human anti-TH-VioR667 (Miltenyi Biotech, 130-131-157, 1:50) to label the dopaminergic neurons. Following multiple washes, the tissues were dehydrated again, incubated in dichloromethane (DCM), and cleared in dibenzyl ether (DBE). For the optimized version of the iDISCO+ protocol, brains were incubated with primary conjugated antibodies for 4 weeks at 37°C, with a top-up incubation on day 7 using the same concentration as the initial dose. α-syn aggregates were stained with the conjugated rabbit anti-pS129 α-syn-Alexa555 (Abcam, ab313137, 1:300), and dopaminergic neurons were labeled using human anti-TH-VioR667 (Miltenyi Biotech, 130-131-157, 1:50). The tissues were subsequently washed and cleared as previously described following dehydration and incubation in DCM and DBE. To test the relevance of the dopamine transporter (DAT) as a dopaminergic cell marker using iDISCO+, we evaluated the rabbit anti-TH antibody (Merck, AB152, 1:500) and the anti-DAT antibody (Merck, MAB369, 1:500) following the original iDISCO+ procedure. The tissues were incubated for 7 days with primary antibodies, followed by a 7-day incubation with the corresponding secondary antibodies (anti-rabbit Cy3, Jackson, 711-165-152, 1:500, and anti-rat Alexa647, Jackson, 712-605-153, 1:500), before the clearing steps were performed.

### LSM

Within 3 days after iDISCO+ clearing, the brains were imaged using an in-house-built LSM, *mesoSPIM* (Center for Microscopy and Image Analysis - ZMB, University of Zurich).^21^ Whole brains were imaged in the horizontal plane using a 4 µm z-step with a Mitutoyo M Plan Apo 2×/0.055 objective (Mitutoyo, Japan), resulting in a 2.75× 2.75×4 µm pixel resolution. A 488 nm excitation laser was used for autofluorescence acquisition, and 561 nm and 647 nm lasers were used for α-syn and TH detection, respectively, using 20 ms of exposure from both sides and each laser, with a quadband 405/488/561/640 nm emission filter, for a total acquisition time of 1 hour (20 minutes per channel). The generated 4 tiles led to a final ∼13.7×24.0×9.3 mm FOV that was subsequently stitched, cropped and fused using Fiji - BigStitcher (NIH, USA), resulting in an approximately 7.1×12.7×6.4 mm FOV. Zoomed-in images of the pS129 α-syn signal were generated using an in-house-built LSM (Benchtop *mesoSPIM)* equipped with a Mitutoyo M Plan Apo 20×/0.28(t3.5) objective, resulting in a 0.2125×0.2125 µm pixel resolution with an FOV of 629×1074 µm.^22^ Two-micrometer z-step z-stacks from the SNc and striatum were acquired at 561 nm, with 20 ms of exposure from the left side (corresponding to the ipsi-injected side) and a quadband 405/488/561/640 nm emission filter. Single slides or 100 µm-thick maximum intensity projection (MIP) were used to show the specificity, consistency and reliability of α-syn pathology labeling and detection in these regions. Images from the 2 mm-thick coronal cleared tissue sections were also acquired with a Benchtop *mesoSPIM* microscope using a Mitutoyo M Plan Apo 5×/0.14 objective for a 0.85×0.85 µm pixel resolution.

### Whole-brain LSM-MRI image coregistration

To achieve precise, robust, and efficient alignment and normalization of LSM and MRI data across multiple modalities with different spatial scales and to correct and assess clearing-induced distortion in the brain regions of interest, we developed a fully automated, coarse-to-fine coregistration framework (**Fig 1**). This framework maps high-resolution LSM and multiparametric MRI data (T1w, QSM/SWI, and DTI maps) into a standardized, 20-μm isotropic common space, specifically DevCCF^52^. This alignment is achieved via two parallel-accelerated cascaded workflows based on diffeomorphic symmetric normalization (SyN)^63^. In the MRI workflow, raw T1W images underwent N4 bias correction and initial coarse alignment using intensity-based center-of-mass translation^51^, followed by multistage registration consisting of rigid, affine, and nonlinear SyN transformations to the DevCCF T2-weighted template. All the stages were constrained by a brain mask and optimized with Mattes mutual information as the similarity metric. For DTI, raw FA maps were first N4 bias-corrected and then registered to the template space using a multistage strategy consisting of rigid, affine, and nonlinear (SyN) transformations, with local cross-correlation as the similarity metric and registration constrained within a brain mask. The resulting deformation fields of the FA maps were propagated to the AD, MD, and RD maps. The SWI and QSM were subsequently warped to the DevCCF space using the T1w-to-template deformation fields.

In the parallel LSM workflow, the raw autofluorescence volume (488 nm) was first downsampled using mean intensity pooling and aligned to the 20 µm DevCCF LSM template using a rigid transformation followed by SyN registration with affine pre-alignment, both driven by Mattes mutual information and constrained by an atlas-derived brain mask. The resulting image was then masked with the same binary mask to suppress background autofluorescence and clearing artifacts. This masked volume served as the moving image for a cross-modal refinement step: after N4 bias correction, it was registered to the template-aligned T1W MRI in DevCCF space via SyN registration to maximize anatomical concordance between modalities. The concatenated deformation fields from both stages (i.e. intra-modal LSM-to-template and inter-modal LSM-to-MRI) were subsequently propagated to the detection channels, warping the pS129 (561-nm) and TH (647-nm) signals into the same space.

### Evaluation of the coregistration accuracy and clearing-induced distortion

To validate the spatial alignment between the coregistered LSM and MRI datasets, we performed a landmark-based assessment using anatomically stable structures. Seven distinct anatomical landmarks were selected on the basis of their unequivocally identifiable and high-contrast boundaries across both modalities^28,33^. These included the ventricular system (lateral and fourth ventricles, including the lateral recess) and major white-matter tracts extending along the rostrocaudal axis (corpus callosum, including the anterior forceps, olfactory and temporal limbs of the anterior commissure, internal capsule, and fimbria/fornix). Coregistration accuracy was quantified by extracting 2D orthogonal slices (sagittal, horizontal, and coronal) at the centroids of these landmarks and measuring the mean spatial displacement and variance between the corresponding boundaries.

To enable a region-specific analysis of tissue deformation induced by the clearing process, the ABA atlas^29^ was back-projected into the individual space of each mouse’s LSM and MRI datasets. This was achieved through an inverse registration pipeline that applied inverse affine and SyN warps into the rigid-aligned native space reconstructed from the raw acquisition data. Clearing-induced tissue distortion was then quantified by comparing the geometric properties of individual brain regions in the LSM space versus the MRI space. For each mapped region, we extracted 3D voxel coordinates and performed eigenvalue decomposition on the coordinate covariance matrix to determine the three principal axes. Deformation was evaluated using three primary metrics: the mean scaling factor, which was calculated as the ratio of the principal axis lengths; the volume retention ratio; and an anisotropy score. The anisotropy score was defined as the coefficient of variation of the scaling factors across the three axes. Finally, to categorize the susceptibility of different brain regions to structural distortion, we performed K-means clustering (k=3) based on the mean scaling factor and anisotropy score. Based on their cluster centroids, regions were classified into three deformation profiles: stable (scaling near 1.0, low anisotropy), uniformly shrunk (reduced volume, low anisotropy), and highly distorted (high anisotropy reflecting severe nonuniform deformation). This quantification of coregistration accuracy and tissue distortion was evaluated using a validation group of three animals (n=3).

## Quantification and statistical analysis of pS129, TH and multiparametric MRI

Whole-brain LSM and multiparametric MRI data from 37 mice (18 PFF-injected and 19 monomer-injected controls) were processed to facilitate high-fidelity, cross-modal integration for voxelwise and regional analyses (**Fig 1**). First, fully automatic anatomical annotation was achieved utilizing a hierarchical parcellation framework encompassing the ABA, the DeCCF atlas, a highly granular molecular atlas for striatal subregions^30^, and a TH distribution-derived consensus atlas for SNc and VTA subregions. This consensus atlas was dynamically generated strictly from the endogenous TH signal of the monomer-injected group. Following the extraction of the largest 3D-connected components, the SNc and VTA were geometrically subdivided into six distinct spatial subregions: lateral posterior-SNc, medial posterior-SNc, medial anterior-SNc, lateral anterior-SNc, anterior-VTA, and posterior-VTA via coordinate-based centroid calculations and ABA-based annotation for the VTA.

To isolate the genuine fluorescence signal from pathological α-syn aggregations and TH+ neuron, we employed an automatic, region-adaptive 3D computational pipeline. Nonspecific background fluorescence was mitigated using a robust random sample consensus (RANSAC) regression algorithm, mathematically modeling the pS129 signal (561 nm) against the tissue autofluorescence (AF) reference channel (488 nm). Following RANSAC correction, localized background subtraction was executed utilizing a 3D Gaussian filter (σ=3.0). Crucially, segmentation thresholds for pS129 were established using a highly stringent empirical percentile (99.99367%, effectively equivalent to a robust 4-sigma threshold) derived strictly from the local background distributions of monomer-injected control mice, guaranteeing the highly specific detection of α-syn pathology. A subsequent 3D distance-transform-based watershed algorithm coupled with strict size-exclusion filtering was used to accurately segment and declump discrete pS129-positive aggregates and calculate their volumetric coverage ratios. Robust local background subtraction (σ=15) and ipsi/contra hemisphere fluorescence intensity normalization were applied to the TH channel to quantify dopaminergic integrity. In parallel, comprehensive regional mean values were extracted for MRI-derived structural and magnetic signatures, encompassing QSM and DTI indices (FA, AD, RD, MD). Group-level statistical comparisons of spatial distributions and signal intensities were evaluated using the nonparametric Mann-Whitney U test. Multimodal correlation analyses were conducted to elucidate the systemic patterns of pathological propagation, and comprehensive 3D spatial covariance networks were constructed by computing pairwise Pearson correlation coefficients across all hierarchical anatomical nodes. To strictly control for false positive topological connections, the resulting network edges were filtered through a rigorous dual-thresholding paradigm: retaining only positive correlations demonstrating high structural covariance (R>0.5) that concurrently survived Benjamini–Hochberg multiple comparisons correction for the false discovery rate (FDR, q<0.01). Significant covariance matrices were subsequently mapped back into the 3D anatomical space and visualized using hierarchical chord diagrams to delineate the intricate macroscopic topology of disease propagation. To map spatiotemporal pathology, multimodal whole-brain alterations were quantified at voxel and regional resolutions using intergroup differences normalized to the 95th percentile. Significant neuroanatomical changes were delineated by overlaying voxelwise statistical maps onto autofluorescence backgrounds utilizing maximum intensity projection (MIP).

## 3D representations

Video of the 3D representative dopaminergic system and α-syn spread throughout the brains of PFF-injected mice was obtained with Imaris (Oxford instruments, UK). Video editing was performed with Adobe express (Adobe, USA).

## DECLARATION

### Ethics approval and consent to participate

Ethics approval: All animal experiments were performed according to the ethical standards of the institution at which the studies were conducted (Regierungspräsidium Giessen, Germany V54-19c2015h01GI 20/28 Nr. G32/2022).

Consent to participate: Not applicable

## Consent for publication

Not applicable

## Authors’ contributions

BC, MH, FG and RN conceived and designed the study. AKR, MTH, and FFG handled the mice, performed all the in vivo experiments and prepared the manual histology quantification. BC performed all the ex vivo experiments and image acquisition. SD wrote the code for automated registration and analysis. BC and FD prepared the figures and videos. BC, SD, MK, SP and RN analyzed the data. PZ and HW analyzed the SWI data. RC, VLD and TMD provided PFFs and guidance on their handling and use. BC, SD and RN wrote the first draft. All the authors contributed to the revision of the manuscript. All the authors read and approved the final manuscript.

## Acknowledgments

The authors would like to thank Dr. Nikita Vladimirov, Dr. Marco Garbelli and Dr. José María Mateos Melero at the URPP AdaBD, Brain Research Institute and the Center for Microscopy and Image Analysis (ZMB), University of Zurich, Dr. Huang Sheng-Fu and Dr. Annika Keller at the Department of Neurosurgery, Zürich University Hospital, and Dr. Hikari Yoshihara at Institute for Biomedical Engineering, ETH Zurich & University of Zurich for technical assistance.

## Data availability

Data are available upon request.

