## supplemental files for "Multiscale brain-wide mapping of α-synuclein-driven dopaminergic degeneration and white matter impairment in α-synucleinopathy mice"

**Supplementary Videos**

[10.5281/zenodo.17770001](https://doi.org/10.5281/zenodo.17770001).

**SVideo 1**: Representative video of classical whole-brain iDISCO+ protocol showing incomplete pS129 α-syn (white) and TH (red) labelling in the striatum 12 wpi of α-syn PFFs in the SNc; scale bar = 1.5 mm. α-syn: α-synuclein; PFFs: preformed fibrils; pS129: phosphorylated serine of α-syn at position 129; SNc: substantia nigra pars compacta; TH: tyrosine hydroxylase; wpi: weeks post injection.

**SVideo 2**: Representative video of enhanced whole-brain iDISCO+ protocol showing complete pS129 α-syn (white) and TH (red) labelling in the brain, 12 wpi α-syn PFFs in the SNc; scale bar = 1.5 mm. α-syn: α-synuclein; PFFs: preformed fibrils; pS129: phosphorylated serine of α-syn at position 129; SNc: substantia nigra pars compacta; TH: tyrosine hydroxylase; wpi: weeks post injection.

**SVideo 3**: 3D representation of enhanced whole-brain iDISCO+ protocol showing complete pS129 α-syn (white) and TH (red) labelling in the brain, 12 wpi α-syn PFFs in the SNc. α-syn: α-synuclein; PFFs: preformed fibrils; pS129: phosphorylated serine of α-syn at position 129; SNc: substantia nigra pars compacta; TH: tyrosine hydroxylase; wpi: weeks post injection.

**SVideo 4**: Representative video of enhanced whole-brain iDISCO+ protocol showing no pS129 α-syn (white) but complete TH (red) labelling in the brain, 12 wpi of α-syn monomers in the SNc; scale bar = 1.5 mm. α-syn: α-synuclein,; pS129: phosphorylated serine of α-syn at position 129; SNc: substantia nigra pars compacta; TH: tyrosine hydroxylase; wpi: weeks post injection.

**Supplementary Analysis Codes and Data**

<https://github.com/dddshzy/PDmice-LSM-MRI-DataFusion>

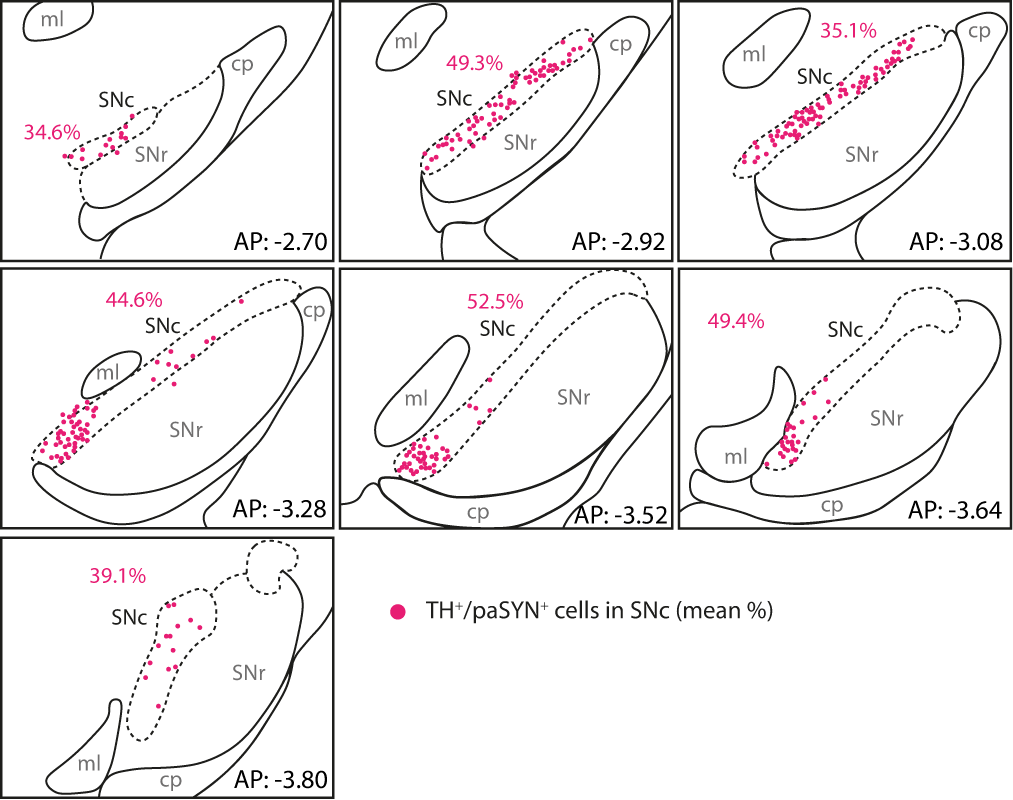

**SFig 1 Unilateral injection site of monomer and PFF in the SNc of mice.** PFFs: preformed fibrils; SNc: substantia nigra pars compacta; SNr: substantia nigra reticulata; TH: tyrosine hydroxylase;

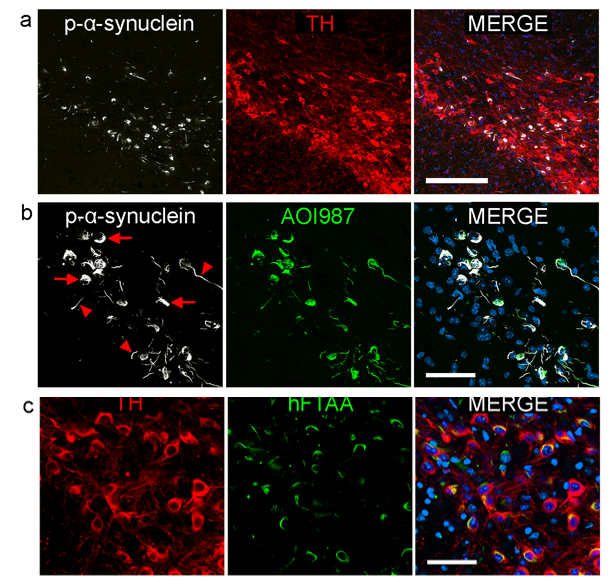

**SFig 2.** **Detection and Amyloid Dye Co-localization of Pathological α-Synuclein in Dopaminergic Neurons of the SNc in the PFF Mouse Model.** (**a**) Representative immunofluorescence images from classical freefloating immunohistochemistry of pS129 α-syn (white), TH (red) and DAPI (blue) in the SNc of PFF-injected mice showing intracellular α-syn aggregates in TH-positive dopaminergic neurons; scale bar = 200 µm. (**b**) and (**c**) AOI987 and hFTAA, which are known to bind β-sheetrich amyloid fibrils, colocalize with the pS129-positive signal in TH+ dopaminergic cells. Red arrows show soma pathology, and red triangles show neurite pathology; scale bars = 50 µm.

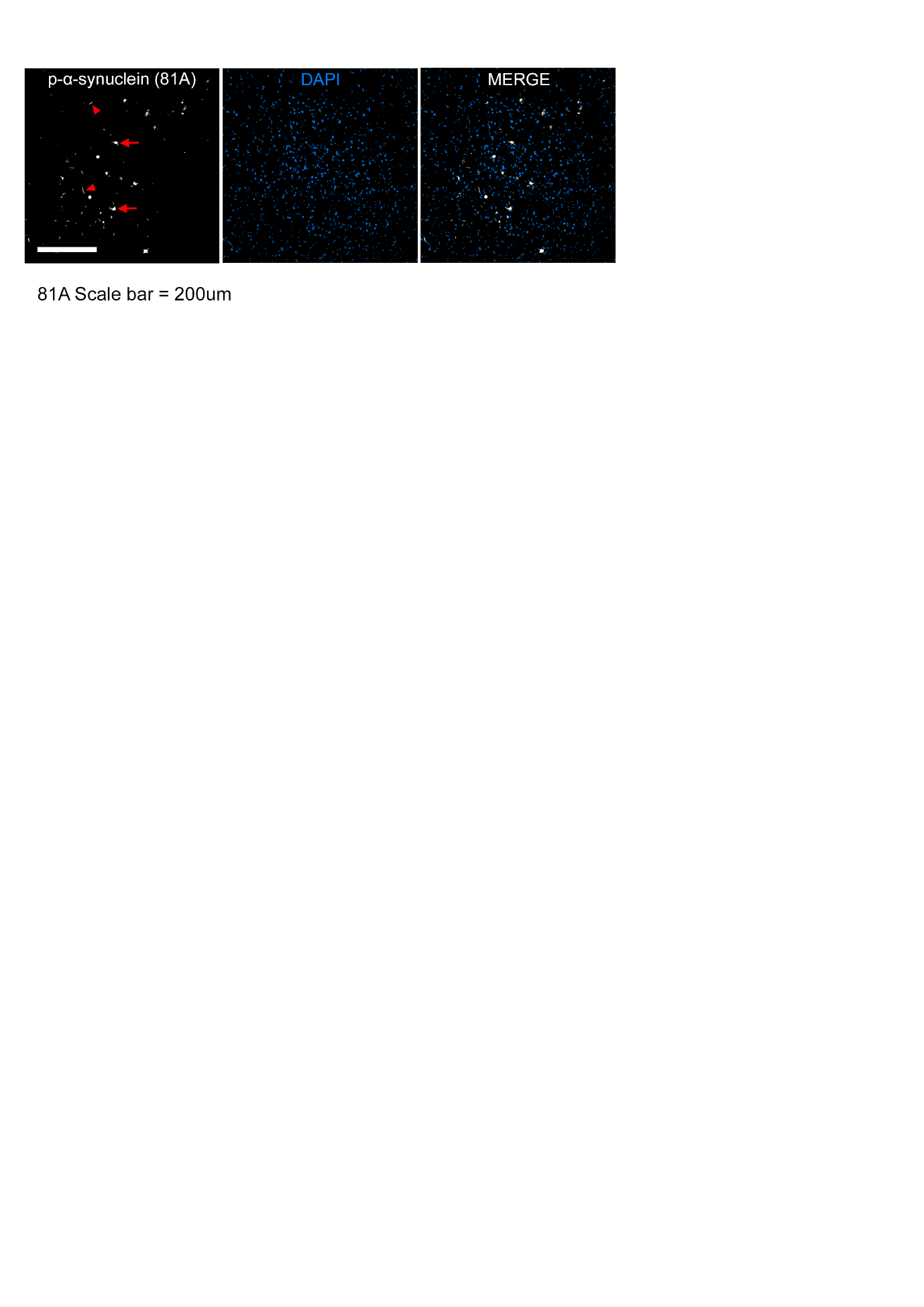

**SFig 3. The 81A clone reproduces staining similar to that of EP1536Y, which targets pS129 α-syn.** Representative immunofluorescence images from classic free-floating sections labeled with 81A clone α-syn pS129 (white) and DAPI (blue) in the PPNs of PFF-injected mice, showing soma (red arrows) and neuritic (red triangles) pathologies; scale bar = 200 µm. α-syn: α-synuclein; PFFs: preformed fibrils; pS129: phosphorylated serine of α-syn at position 129; PPN: pedunculopontine nucleus.

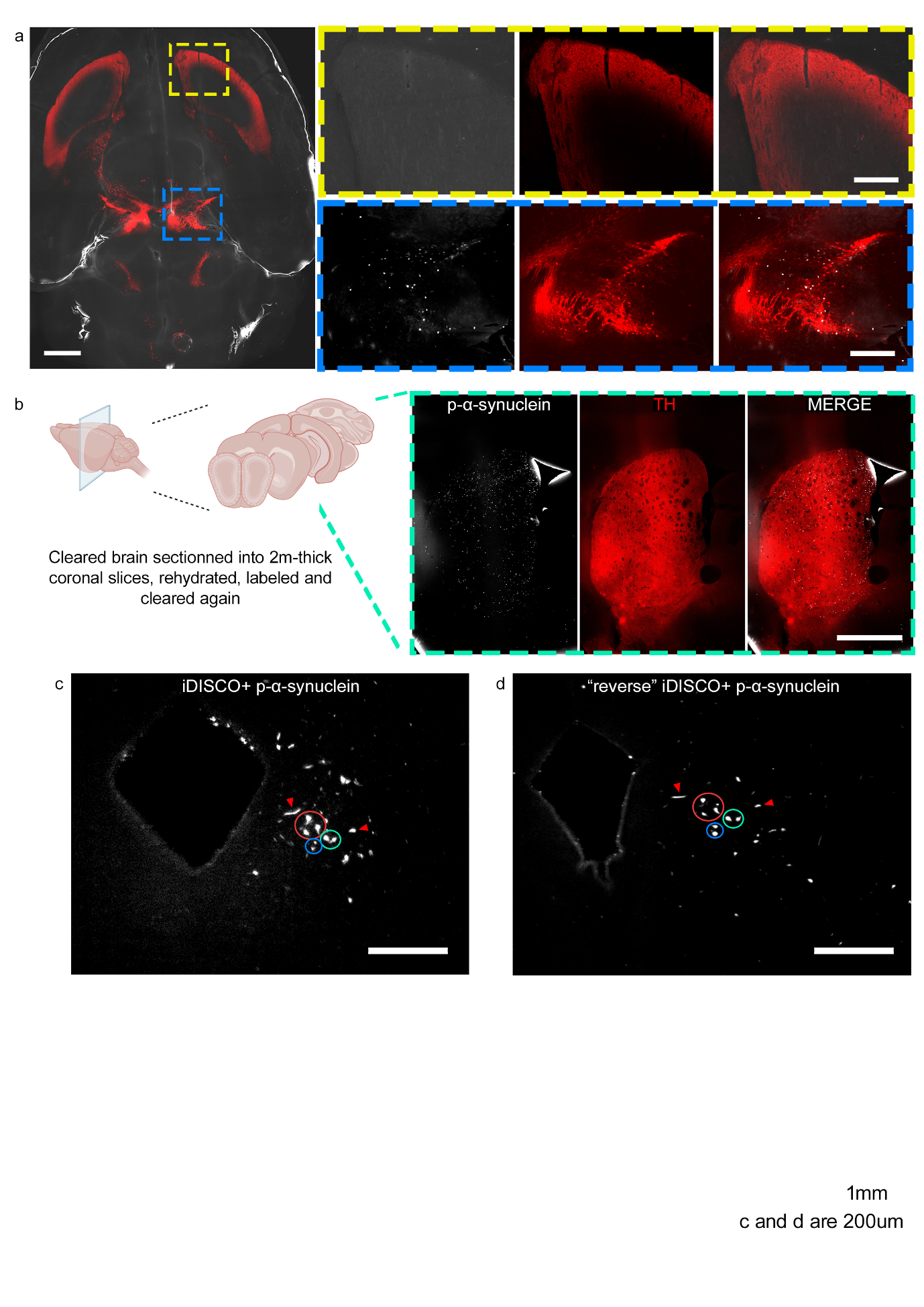

**SFig 4. Rehydration, relabeling and reclearing of 2 mm-thick coronal sections after the iDISCO+ protocol revealed specific pS129- and TH-positive signals in the striatum.** a) Representative whole-brain iDISCO+ slices showing partial α-syn pS129 (white) and TH (red) labelling in the striatum (yellow box), whereas the SNc (blue box) displays a robust signal 12 wpi of PFFs into the SNc; scale bars = 1.5 mm (overview) and 400 µm (digital zoom, post-capture enlargement). b) Green dotted rectangles illustrate 2 mm thick coronal sections obtained after the cleared and previously incompletely stained brains were cut. In the right panel, pS129- and TH-positive signals are shown in the striatum after rehydration, relabeling and reclearing the same mouse tissue, so-called “reverse iDISCO+”; scale bar = 1 mm. c and d) Representative coronal slices of iDISCO+ (c) and reverse iDISCO+ (d) showing the identification of the same α-syn aggregates in the PAG, a properly stained region with classical iDISCO+; scale bars = 200 µm. α-syn: α-synuclein; PAG: periaqueductal gray; PFFs: preformed fibrils; pS129: phosphorylated serine of α-syn at position 129; SNc: substantia nigra pars compacta; TH: tyrosine hydroxylase; wpi: weeks post injection.

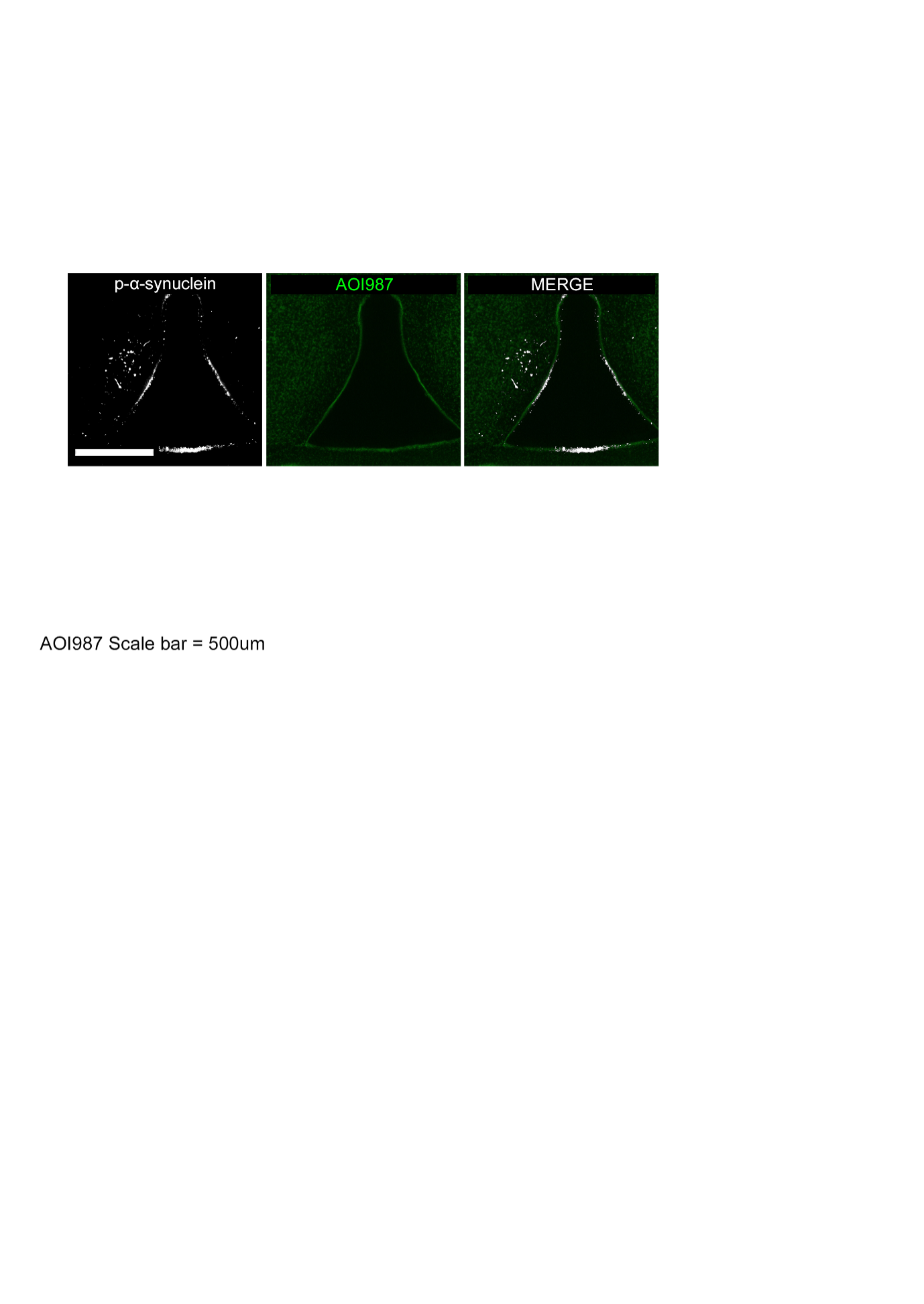

**SFig 5. AOI987 does not detect pS129 α-synuclein using the iDISCO+ protocol.** Representative local horizontal iDISCO+ image showing localized α-syn pS129 (white) signal in the PPN, which does not colocalize with AOI987 labeling (green); scale bar = 500 µm. α-syn: α-synuclein; pS129: phosphorylation of α-synuclein at serine 129; PPN: pedunculopontine nucleus.

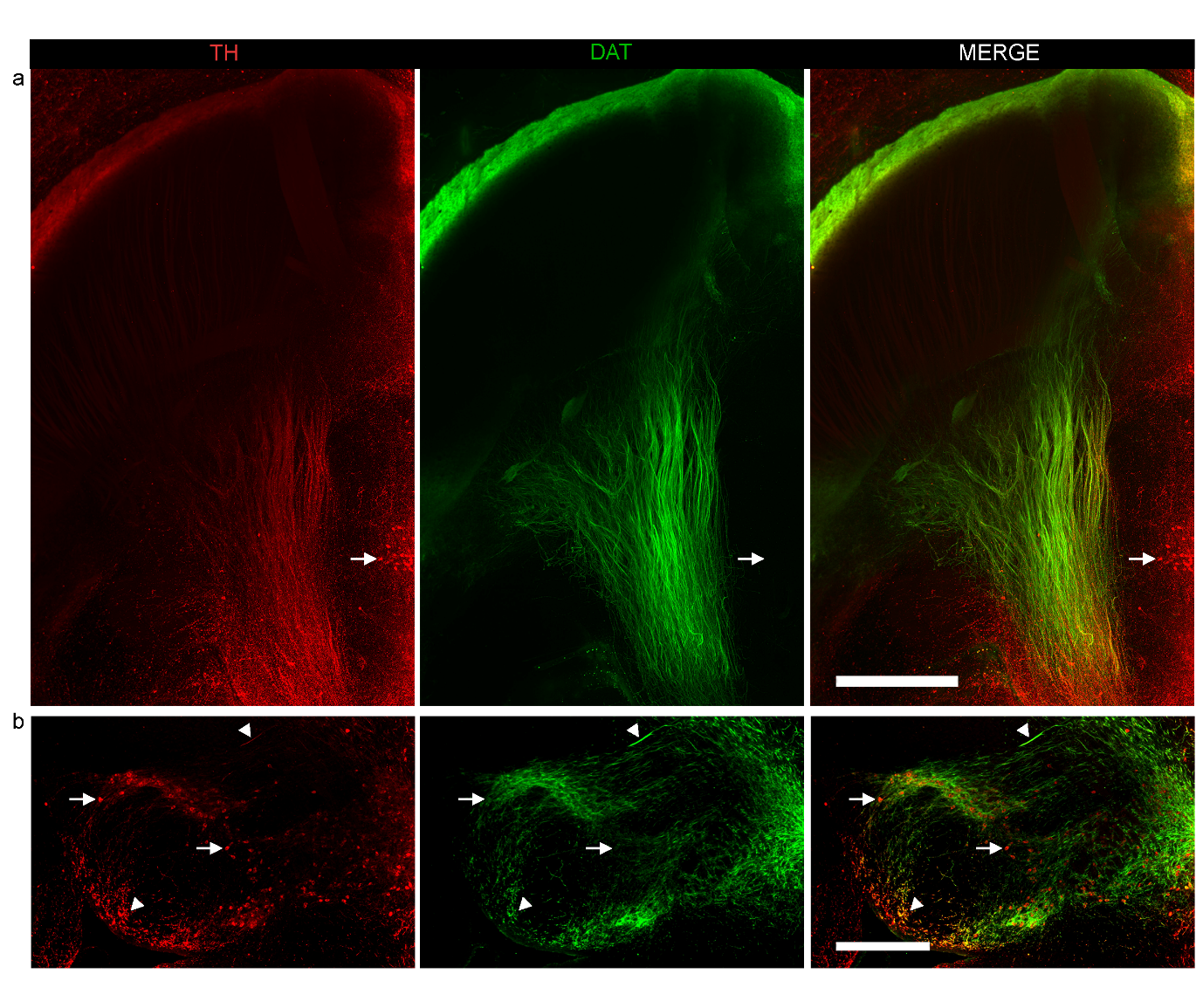

**SFig 6 Identification of DAT-negative but TH-positive neuronal populations in the mouse brain.** a) Representative horizontal slice labeled using the classical unoptimized iDISCO+ protocol, showing regions of TH and DAT colocalization within the MFB and the outer striatum. The white arrow indicates TH-positive but DAT-negative cell bodies located in the PVH; scale bar = 800 µm. b) A similar pattern is observed in the SNc and closed regions, where some neuronal somas are TH-positive and DAT-negative (white arrows), whereas axonal processes show strong colocalization of TH and DAT (white triangles), notably in the SNr (bottom white triangle); scale bar = 500 µm. α-syn: α-synuclein, DAT: dopaminergic transporter, MFB: medial forebrain bundle, SNc: substantia nigra pars compacta, SNr: substantia nigra reticulata, PVH: paraventricular hypothalamic nucleus, TH: tyrosine hydroxylase

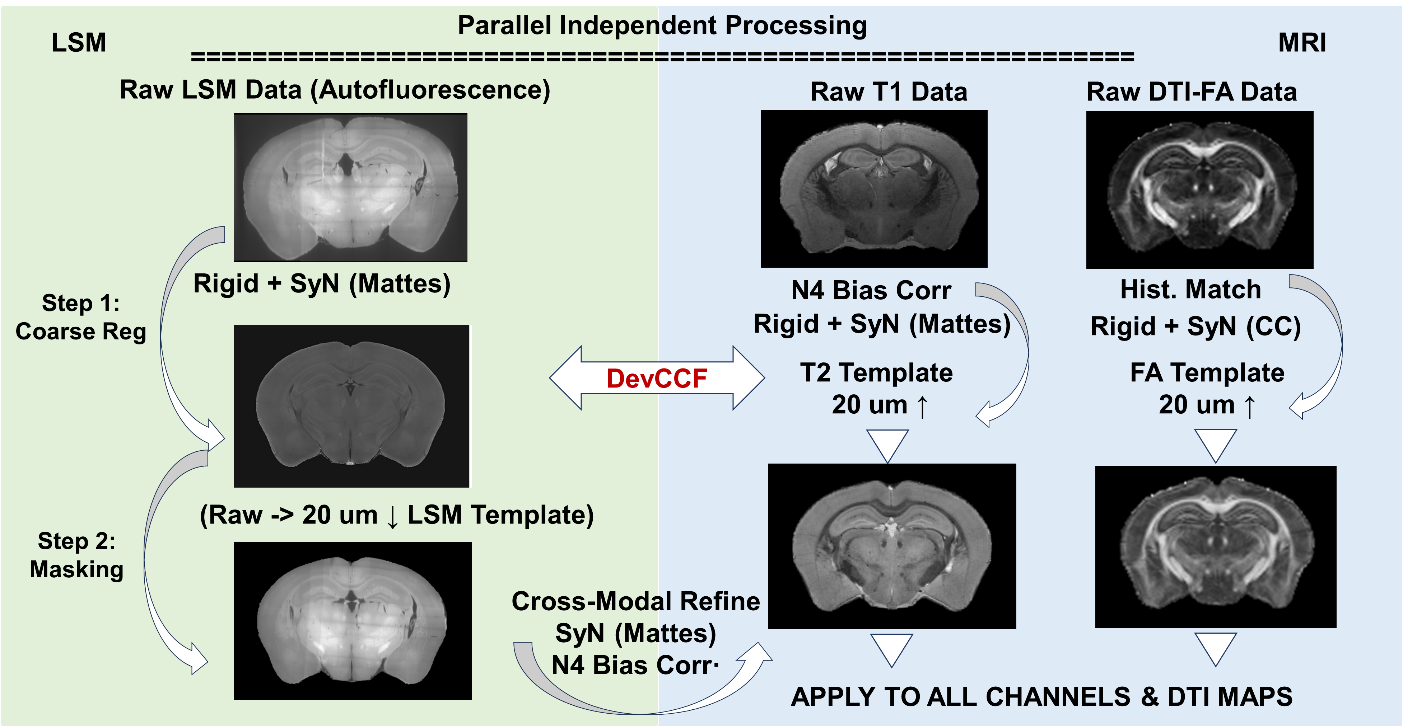

whole-brain LSM/MRI volume ratio

- CR2403: 73.76%
- CR2421: 74.02%
- CR2413: 75.95%

**SFig 7**: **Schematic diagram of the LSM-MRI Registration Pipeline.** The implemented pipeline employs a parallelized, multi-stage registration framework based on the ANTs (Advanced Normalization Tools) library to align LSM and MRI data within a unified 20 micrometers coordinate space. The pipeline consists of two concurrent processing streams. In the MRI workflow, raw T1-weighted images are pre-processed with N4 bias field correction and registered to a standard T2 template (upsampled to 20 micrometers) using a composite Rigid and SyN transformation with Mattes mutual information. Separately, Diffusion Tensor Imaging (DTI) maps are processed by registering Fractional Anisotropy (FA) to an FA template via Cross-Correlation (CC) with Histogram Matching; the resulting transformation is applied to associated diffusivity maps (AD, MD, RD). In parallel, the LSM workflow performs sequential steps: raw LSM (488nm) is geometrically corrected and registered to an LSM template using Rigid and SyN, followed by custom masking to remove noise and artifacts. Finally, the masked volume acts as the moving image and is refined by registering it to the aligned MRI T1 volume, incorporating N4 correction as a pre-processing step to ensure robust alignment. The transformations in LSM workflow are propagated to 561 (pS129) and 640 nm (TH).

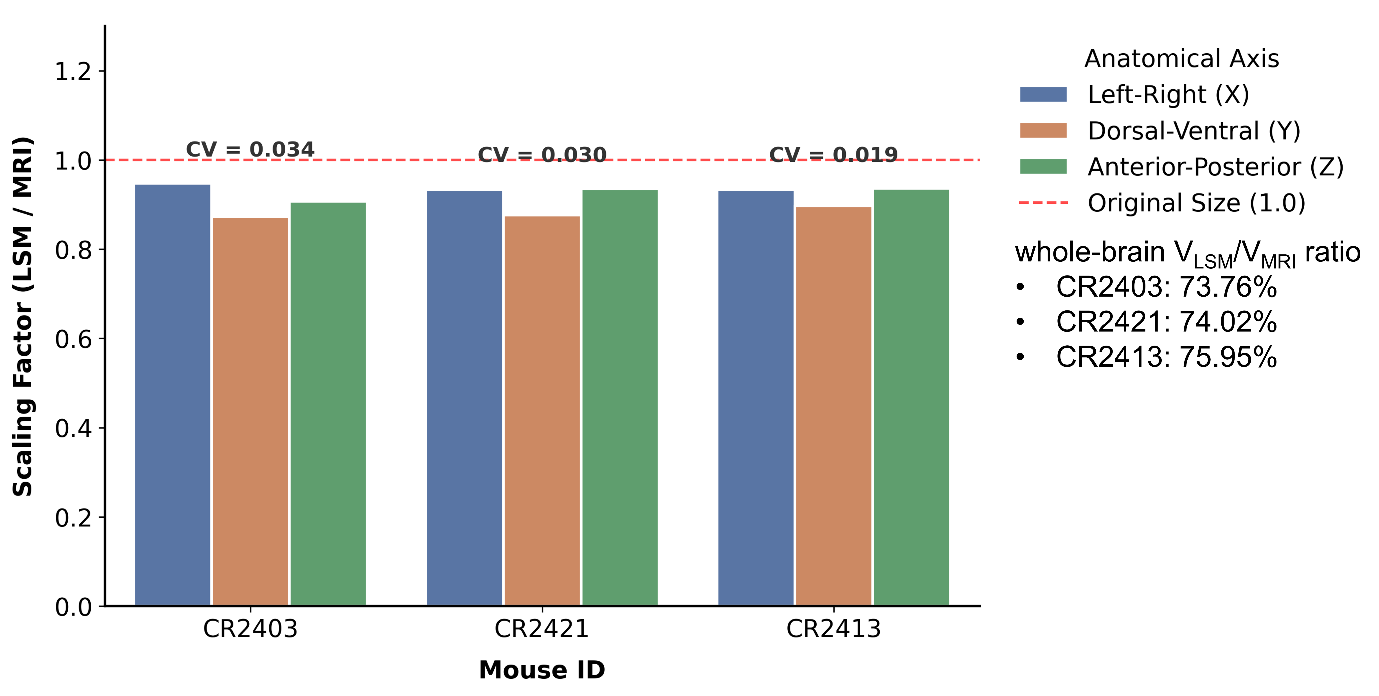

**SFig 8.** **Global deformation analysis and anisotropic scaling assessment following tissue clearing.** Quantitative evaluation of global tissue shrinkage across the three principal macroscopic axes (X, Y, and Z). The overall mean scaling factor across all subjects is approximately 0.91, indicating a consistent global tissue shrinkage of ~9% induced by the clearing protocol. Notably, the tissue deformation exhibits a distinct directional bias: while the X and Z axes remain relatively preserved (scaling factor > 0.93), the Y-axis (dorsal-ventral direction) consistently undergoes the highest degree of compression (scaling factor < 0.90). High inter-subject consistency is confirmed by the exceptionally low anisotropy coefficient of variation (CV < 0.035), demonstrating that this anisotropic deformation pattern is highly stable and reproducible across different biological replicates.

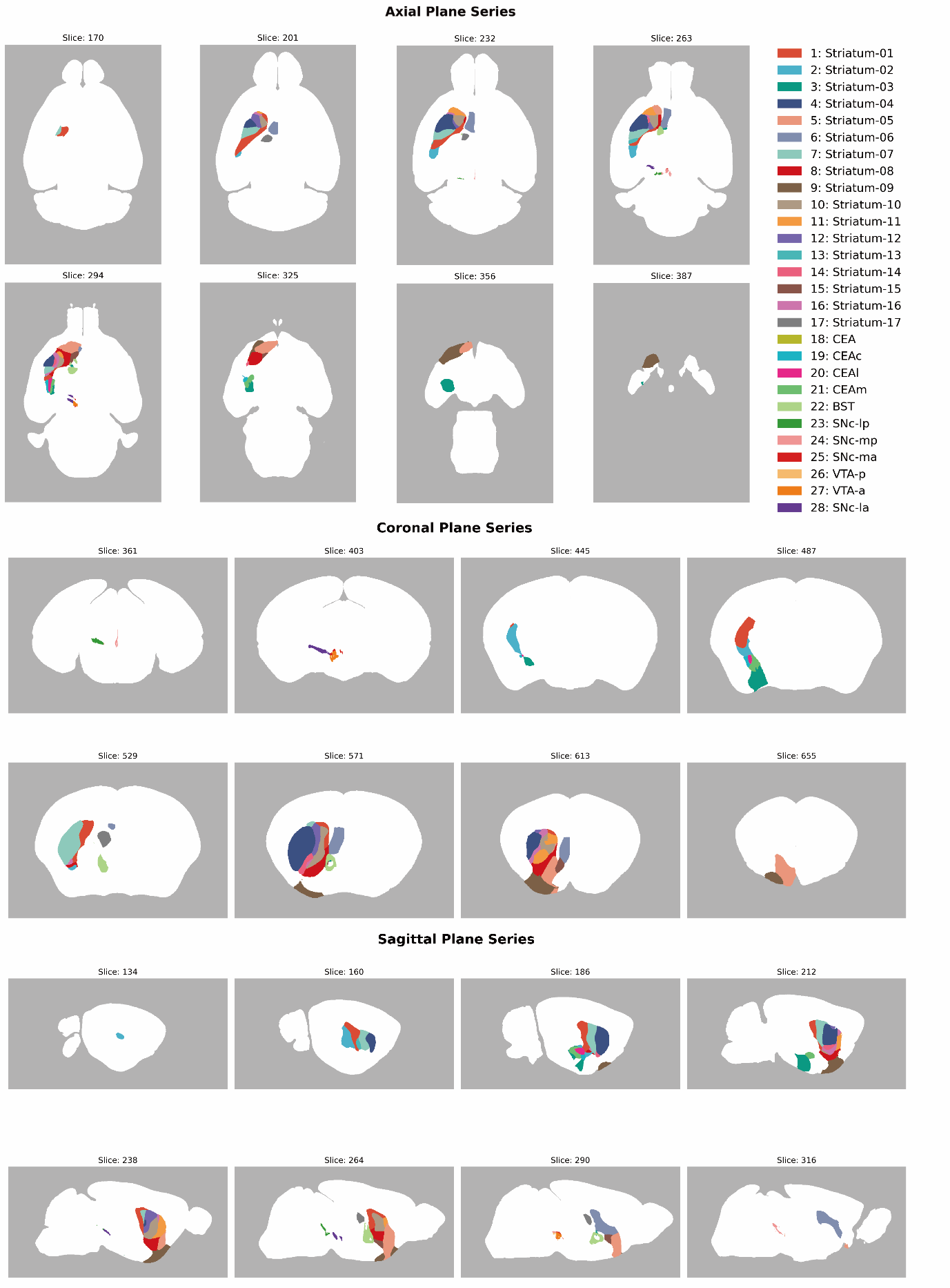

**SFig 9. Multi-planar mapping of targeted neuroanatomical regions of interest (ROIs).** Representative 2D slice series across the sagittal, axial, and coronal planes illustrating the spatial distribution and boundaries of the selected ROIs within the ipsilateral (left) hemisphere. The distinctly color-coded ROIs are aggregated from three distinct hierarchical parcellation frameworks: a high-resolution Molecular Atlas (defining 17 striatal sub-regions), the standard ABA (defining the CEA and BST), and a dynamically generated custom consensus atlas (geometrically subdividing the SNc and VTA). ABA: Allen Brain Atlas, BST: Bed nuclei of the stria terminalis, CEA: Central amygdalar nucleus, CEAc: Central amygdalar nucleus, capsular part, CEAl: Central amygdalar nucleus, lateral part, CEAm: Central amygdalar nucleus, medial part, ROI: Region of interest, SNc: Substantia nigra pars compacta, SNc-la: Substantia nigra pars compacta, lateral anterior, SNc-lp: Substantia nigra pars compacta, lateral posterior, SNc-ma: Substantia nigra pars compacta, medial anterior, SNc-mp: Substantia nigra pars compacta, medial posterior, VTA-a: Ventral tegmental area, anterior, VTA-p: Ventral tegmental area, posterior.

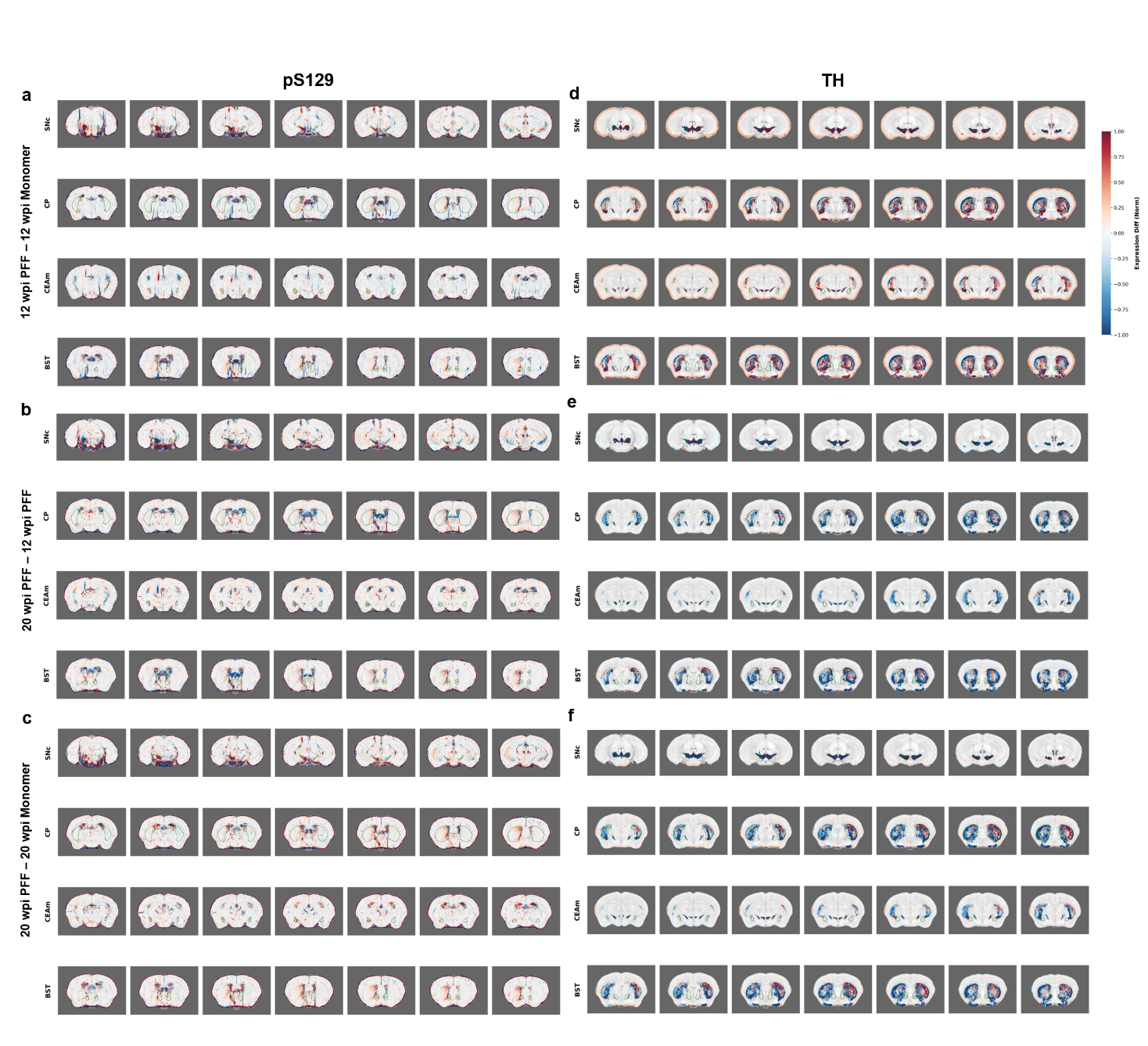

**SFig 10. Voxel-wise spatial distribution of expression differences across key neuroanatomical ROIs.** Representative 2D multi-slice series illustrating the normalized spatial expression differences of targeted molecular markers (pS129 and TH) across varying experimental cohorts. pS129 fluorescence intensity (FI) differences for **(a)** 12 wpi PFF vs. 12 wpi Monomer, **(b)** 20 wpi PFF vs. 12 wpi PFF, and **(c)** 20 wpi PFF vs. 20 wpi Monomer. TH FI differences for **(d)** 12 wpi PFF vs. 12 wpi Monomer, **(e)** 20 wpi PFF vs. 12 wpi PFF, and **(f)** 20 wpi PFF vs. 20 wpi Monomer. Differential maps are presented as maximum absolute projections computed over a 10-slice rolling window and are seamlessly overlaid onto a mean tissue autofluorescence (AF) template. The divergent color scale indicates bidirectional relative expression changes normalized to the 95th percentile. Specific targeted sub-regions are delineated by dashed dark green contours to provide anatomical localization. AF: Autofluorescence, BST: Bed nuclei of the stria terminalis, CEAm: Central amygdalar nucleus, medial part, CP: Caudoputamen, FI, fluorescent intensity, PFF: Preformed fibrils, ROI: Region of interest, SNc: Substantia nigra pars compacta, TH: Tyrosine hydroxylase, wpi: Weeks post-injection.

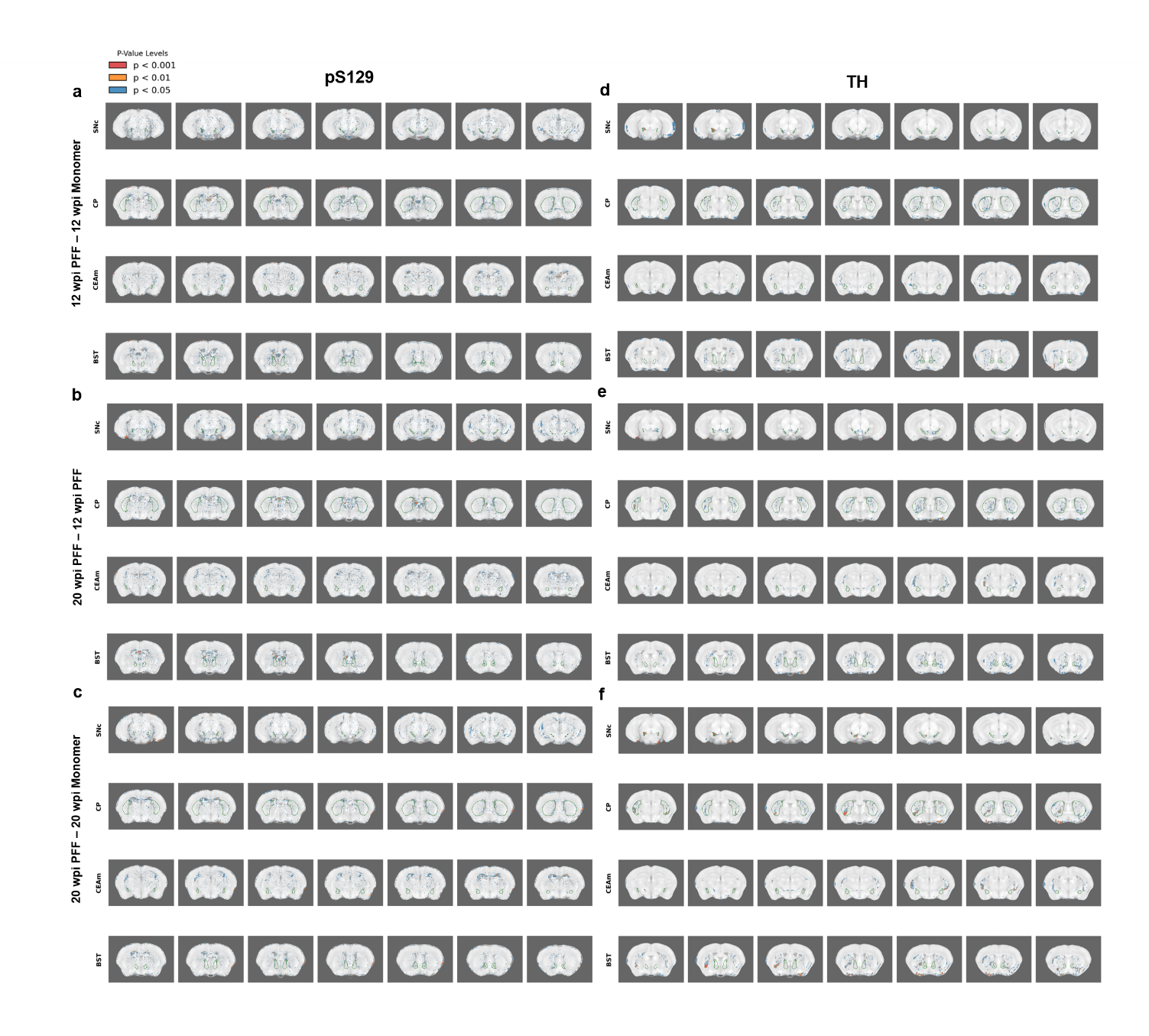

**SFig 11. Voxel-wise statistical significance mapping of targeted molecular markers across key neuroanatomical ROIs.** p-value maps of pS129 fluorescence intensity (FI) differences for **(a)** 12 wpi PFF vs. 12 wpi Monomer, **(b)** 20 wpi PFF vs. 12 wpi PFF, and **(c)** 20 wpi PFF vs. 20 wpi Monomer, and of TH FI differences for **(d)** 12 wpi PFF vs. 12 wpi Monomer, **(e)** 20 wpi PFF vs. 12 wpi PFF, and **(f)** 20 wpi PFF vs. 20 wpi Monomer. Corresponding representative spatial maps displaying the statistical significance (P-values, derived from two-tailed unpaired Student's t-tests) for the comparative analyses detailed in SFig 12. Images represent the minimum P-value projection across a 10-slice spatial window, superimposed on the mean autofluorescence (AF) background template to contextualize statistical findings within the anatomical framework. Statistical confidence levels are hierarchically color-coded: blue indicates 0.01 ≤ p < 0.05, orange indicates 0.001 ≤ p < 0.01, and red highlights highly significant structural voxels with p < 0.001. The defined boundaries of the examined neuroanatomical structures are consistently outlined by dashed dark green contours. AF: Autofluorescence, BST: Bed nuclei of the stria terminalis, CEAm: Central amygdalar nucleus, medial part, CP: Caudoputamen, PFF: Preformed fibrils, ROI: Region of interest, SNc: Substantia nigra pars compacta, TH: Tyrosine hydroxylase, wpi: Weeks post-injection.

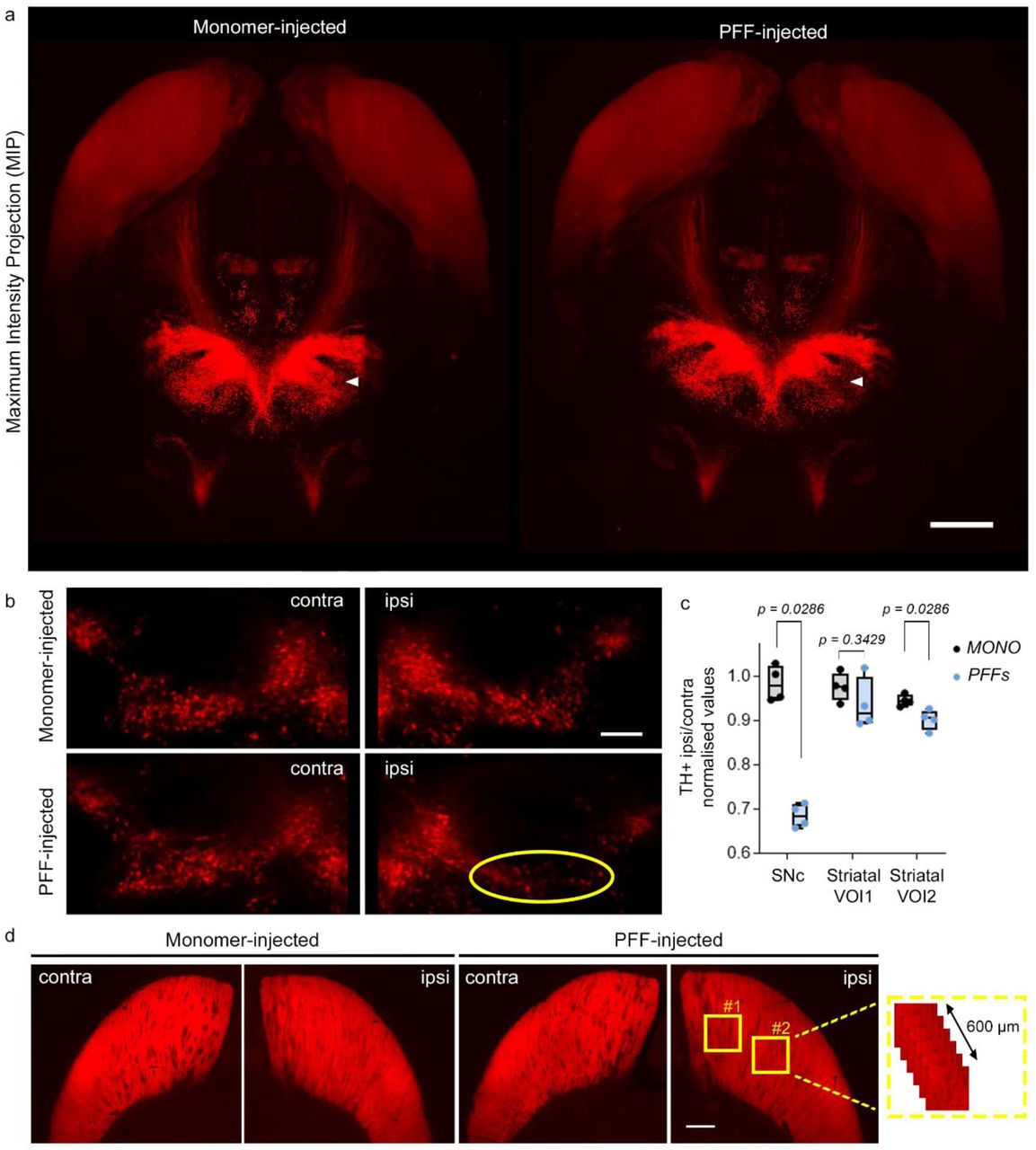

**SFig 12 3D quantification revealed dopaminergic degeneration in the nigrostriatal pathway in PFF-SNc injected mice.** a) TH+ signal MIP representation in monomer- and PFFinjected brains after 12 weeks (the olfactory bulb was masked); scale bar = 1 cm. b) Representative images showing neurodegeneration of dopaminergic SNc neurons (yellow dotted circle) in PFFs injected mice; scale bar = 200 μm. c) Plots showing the ipsi/contra normalized number of TH+ neurons in the SNc and the quantification of TH expression in two different regions of the striatum (box plots represent median and interquartile range, whiskers min/max value; MONO (N=4), PFF (N=4), Mann-Whitney-U test). d) Images showing TH expression in the striatum and 3D quantification method. Yellow squares indicate measured VOIs; scale bar = 500 μm. α-syn: α-synuclein, MIP: maximum intensity projection, MONO: monomers, PFFs: preformed fibrils, SD: standard deviation, SNc: substantia nigra pars compacta, TH: tyrosine hydroxylase, VOI: volume of interest, wpi: weeks post injection.

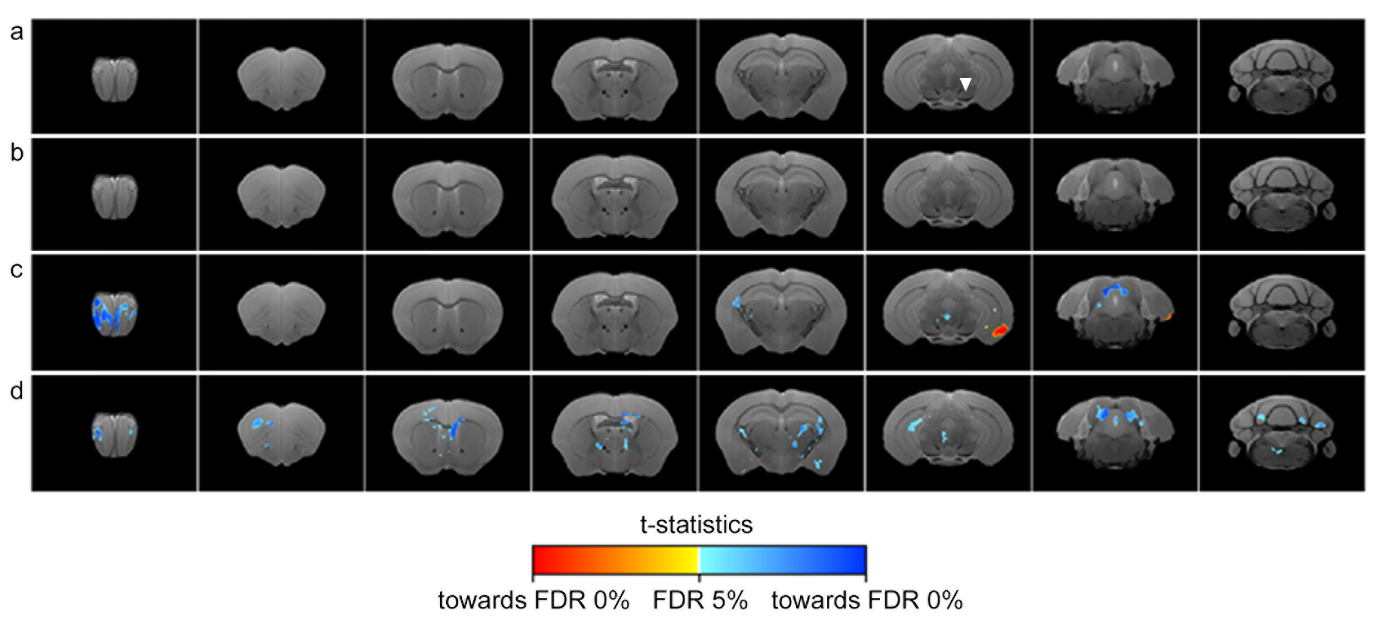

**SFig 13.** **T1-weighted deformation-based morphometry (DBM) results of α-syn PFF or MONO-injected mice at 12 or 20wpi at 60 μm isotropic resolution.** Voxel-wise DBM results are shown as *t*-statistic maps overlaid on the anatomical template: (**a**) 12wpi MONO vs. 12wpi PFF, (**b**) 20wpi MONO vs. 20wpi PFF, (**c**) 12wpi MONO vs. 20wpi MONO, and (**d**) 12wpi PFF vs. 20wpi PFF. Red/yellow colors indicate regional expansion, whereas blue colors indicate regional atrophy. Color bar represents *t*-statistics, thresholded using FDR correction (5%). MONO, α-syn monomers; PFF, α-syn preformed fibrils; wpi, weeks post-injection.

**STable 1. Antibodies, chemicals, and materials used.**

| Technique | Reagent | Dilution  (final concentration) | Source | Identifier |
| --- | --- | --- | --- | --- |
| IHC, IF | Anti-pS129 α-syn, [EP1536Y] (rabbit) | 1:1000 (2.62 µg/ml) | Abcam | ab51253 |
|  | Anti-TH, [CL3049] (mouse) | 1:500 (1 µg/ml) | Merck Millipore | AMAB91112 |
|  | Anti-pS129 α-syn, [81A] (mouse) | 1:1000 (1 µg/ml) | Merck Millipore | MABN826 |
|  | Antibody anti-rabbit – Alexa488 (donkey) | 1:500 (3 µg/ml) | Jackson ImmunoResearch | 711-545-152 |
|  | Antibody anti-mouse – Cy3 (donkey) | 1:500 (3 µg/ml) | Jackson ImmunoResearch | 715-165-150 |
|  | AOI987 | (20µM) | Santa Cruz Biotech | sc-481117 |
|  | hFTAA | (30µM) | Provided by Prof. Peter R. Nilsson |  |
|  | NDS | 3 and 5% | Interchim | UP77719A K |
|  | Triton X-100 | 0.3 and 0.5% | Merck | RES3103T-A101X |
|  | DAPI | 1:1000 (1 µg/ml) | Invitrogen | D1306 |
|  | ProLong Diamond Antifade Mountant |  | Invitrogen | P36970 |
| iDISCO+ | Anti-pS129 α-syn, [EP1536Y] (rabbit) | 1:500 (5.24 µg/ml) | Abcam | ab51253 |
|  | Anti-TH (rabbit) | 1:500 | Merck | AB152 |
|  | Anti-DAT, [DAT-Nt] (rat) | 1:500 | Merck | MAB369 |
|  | Anti-rabbit – Cy3 (donkey) | 1:500 (3 µg/ml) | Jackson ImmunoResearch | 711-165-152 |
|  | Anti-rat – Alexa647 (donkey) | 1:500 (3 µg/ml) | Jackson ImmunoResearch | 712-605-153 |
|  | Anti-pS129 α-syn conjugated with Alexa555, EP1536Y (rabbit) | 1:300  (1.66 µg/ml *2 for top-up at D7) | Abcam | ab313137 |
|  | Anti-TH conjugated with VioR667, [REA1159] (human) | 1:50 (5 µg/ml *2 for top-up at D7 for optimized protocol) | Miltenyi Biotec | 130-131-157 |
|  | AOI987 | (15µM) | Santa Cruz Biotech | sc-481117 |
|  | hFTAA | (5µM) | Provided by Prof. Peter R. Nilsson |  |
|  | Gelatin | 6% | Roth | 4582.3 |
|  | Heparin | 10 µg/ml | Merck | H3393 |
|  | Glycine | 2.3% | Merck | G7126 |
|  | Triton X-100 | 0.3 and 0.5% | Merck | RES3103T-A101X |
|  | 30% H2O2 | 1:5 (v:v), 5% final | Merck | 1072090500 |
|  | DMSO | 20% | Merck | D5879 |
|  | Tween-20 | 0.2% | Merck | P9416 |
|  | Methanol |  | Merck | 179337 |
|  | DCM |  | Honeywell Research Chemicals | 32222 |
|  | DBE |  | Merck | 108014 |

α-syn: alpha-synuclein; D7: day 7; DAPI: 4',6-diamidino-2-phenylindole; DBE: dibenzylether; DCM: dichloromethane; DMSO: dimethyl sulfoxide; H_2_O2: hydrogen peroxide; hFTAA: heptamer formyl thiophene acetic acid; IHC, immunohistochemistry; IF, immunofluorescence; NDS: normal donkey serum; TH: tyrosine hydroxylase.

**STable 2** **Summary of voxel-wise deformation-based morphometry comparisons using permutation-based testing.**

| Comparison | Changes | Test | Significance | Significance Max |
| --- | --- | --- | --- | --- |
| 12wpi MONO vs. 12wpi PFF | Expansion | t-test | False | 0.8768 |
|  | Atrophy | t-test | False | 0.5802 |
| 20wpi MONO vs. 20wpi PFF | Expansion | t-test | False | 0.1424 |
|  | Atrophy | t-test | False | 0.8642 |
| 12wpi MONO vs. 20wpi MONO | **Expansion** | **t-test** | **True** | **0.9976** |
|  | **Atrophy** | **t-test** | **True** | **0.9944** |
| 12wpi PFF vs. 20wpi PFF | Expansion | t-test | False | 0.2742 |
|  | **Atrophy** | **t-test** | **True** | **0.9864** |

Voxel-wise group comparisons of T1-weighted DBM measures were performed using permutation-based *t*-tests with TFCE. Comparisons include α-syn MONO- and PFF-injected mice at 12 and 20 wpi, as well as longitudinal within-group comparisons. Results are reported separately for regional expansion and atrophy. “Significance Max” corresponds to the maximum TFCE-corrected statistic (1 − *p*) across the brain, with values > 0.95 considered statistically significant (TFCE-corrected *p* < 0.05). Significant difference were observed for both expansion and atrophy in MONO-injected mice at 12 vs. 20 wpi, and for regional atrophy in PFF-injected mice at 12 wpi vs. 20 wpi, while no other comparisons reached statistical significance after correction. DBM, deformation-based morphometry; MONO, α-synuclein monomers; PFF, α-synuclein preformed fibrils; TFCE, threshold-free cluster enhancement; wpi, weeks post-injection.

**STable 3:** **Summary of voxel-wise diffusion metric comparisons using permutation-based testing.**

| Comparison | Changes | Metrics | Significance | Significance Max |
| --- | --- | --- | --- | --- |
| 12wpi MONO vs. 12wpi PFF | Increase | AD | False | 0.6228 |
|  | Decrease | AD | False | 0.4574 |
| 20wpi MONO vs. 20wpi PFF | Increase | AD | False | 0.9220 |
|  | Decrease | AD | False | 0.2258 |
| 12wpi MONO vs. 20wpi MONO | Increase | AD | False | 0.9146 |
|  | Decrease | AD | False | 0.6376 |
| 12wpi PFF vs. 20wpi PFF | **Increase** | **AD** | **True** | **0.9608** |
|  | Decrease | AD | False | 0.0844 |
| 12wpi MONO vs. 12wpi PFF | Increase | MD | False | 0.6132 |
|  | Decrease | MD | False | 0.2998 |
| 20wpi MONO vs. 20wpi PFF | Increase | MD | False | 0.9324 |
|  | Decrease | MD | False | 0.2898 |
| 12wpi MONO vs. 20wpi MONO | Increase | MD | False | 0.9156 |
|  | Decrease | MD | False | 0.8526 |
| 12wpi PFF vs. 20wpi PFF | **Increase** | **MD** | **True** | **0.9552** |
|  | Decrease | MD | False | 0.1562 |
| 12wpi MONO vs. 12wpi PFF | Increase | RD | False | 0.5726 |
|  | Decrease | RD | False | 0.1956 |
| 20wpi MONO vs. 20wpi PFF | Increase | RD | False | 0.9222 |
|  | Decrease | RD | False | 0.2424 |
| 12wpi MONO vs. 20wpi MONO | Increase | RD | False | 0.9240 |
|  | Decrease | RD | False | 0.9276 |
| 12wpi PFF vs. 20wpi PFF | Increase | RD | False | 0.9474 |
|  | Decrease | RD | False | 0.3094 |
| 12wpi MONO vs. 12wpi PFF | Increase | FA | False | 0.3120 |
|  | Decrease | FA | False | 0.8676 |
| 20wpi MONO vs. 20wpi PFF | Increase | FA | False | 0.1444 |
|  | Decrease | FA | False | 0.6762 |
| 12wpi MONO vs. 20wpi MONO | Increase | FA | False | 0.9038 |
|  | Decrease | FA | False | 0.7810 |
| 12wpi PFF vs. 20wpi PFF | Increase | FA | False | 0.6540 |
|  | Decrease | FA | False | 0.8322 |

Voxel-wise group comparisons of DTI metrics were performed using permutation-based t-tests with threshold-free cluster enhancement (TFCE). Comparisons include α-syn MONO- and PFF-injected mice at 12 and 20 wpi, as well as longitudinal within-group comparisons. Results are reported separately for increases and decreases in each metric. “Significance Max” corresponds to the maximum TFCE-corrected statistic (1 − *p*) across the brain, with values > 0.95 considered statistically significant (TFCE-corrected *p* < 0.05). Significant effects were observed for increased AD and MD in PFF-injected mice at 12 vs. 20 wpi, while no other comparisons reached statistical significance after correction. AD, axial diffusivity; DTI, diffusion tensor imaging; FA, fractional anisotropy; MD, mean diffusivity; MONO, α-syn monomers; PFF, α-syn preformed fibrils; RD, radial diffusivity; TFCE, threshold-free cluster enhancement; wpi, weeks post-injection.

**Supplemental Methods**

**Preparation of purified α-syn PFFs and α-syn monomers**

Mouse full-length α-syn PFFs were prepared at Johns Hopkins University School of Medicine, Baltimore, USA, as previously described ^1-3^. Briefly, to generate α-syn PFFs, purified full-length mouse α-syn monomers were continuously agitated in a thermomixer (Eppendorf) at 1,000 rpm and 37°C for 7 days. Thereafter, the formed α-syn aggregates were sonicated for 30 s at 20% amplitude with a Branson Digital Sonifier (Danbury, CT, USA). Monomeric α-syn and PFFs were frozen at -80°C. A subset of the stock solutions was then used for quality control assessments. Finally, the aliquoted stock solutions were shipped to Philipps University Marburg (Germany) on dry ice and stored at -80°C. PFFs were thawed, and sterile phosphate-buffered saline (PBS) was added to the solution to achieve a final protein concentration of 2.5 µg/µL. Afterward, the PFFs were sonicated for 90 seconds at 10% amplitude, aliquoted and finally stored at -80°C. On the day of injection, the aliquoted PFFs were thawed and briefly vortexed before injection.

**Free-floating immunolabeling and microscopy**

Coronal sections were rinsed three times in 0.1 M PBS for 10 min, followed by a 1-h incubation in 0.1 M PBS, 5% v/v normal donkey serum (NDS) and 0.5% v/v Triton-X for blocking and permeabilization. Primary antibodies raised against pS129 α-syn (Abcam, ab51253, 1:1000) and tyrosine hydroxylase (TH, Sigma, AMAb 91112, 1:500) were incubated overnight at 4°C in 0.1 M PBS, 3% v/v NDS and 0.3% v/v Tri­ton-X. The sections were then rinsed three times for 10 min each in 0.1 M PBS before being incubated for 2 h with species-specific secondary antibodies (anti-rabbit Alexa488 Jackson, 711-545-152, 1:500 and anti-mouse Cy3 Jackson, 715-165-150, 1:500) suspended in 0.1 M PBS with 3% v/v NDS and 0.3% v/v Triton-X. The sections were then counterstained with 4’,6-diamidino-2-phenylindole (DAPI) and mounted on microscope slides in Prolong Diamond antifade mounting medium. The benzothiazole derivative AOI987 (Santa Cruz Biotech, sc-481117) and the luminescent conjugated oligothiophene hFTAA (heptamer formyl thiophene acetic acid, provided by Prof. Peter R. Nilsson) were tested for their ability to stain α-syn, as they selectively stain protein aggregates by binding to β-sheet-rich amyloid fibrils. ^4,5^. In these cases, the sections were incubated for 30 min in either AOI987 (15 µM) or hFTAA (5 µM) between the secondary antibody and DAPI staining (details of the antibodies are provided in **Supplemental Table 1**). Representative images of the pS129 α-syn-positive aggregates and TH+ cells were obtained at 20× magnification using a TCS SP8 confocal laser scanning microscope (Leica, Germany). The same confocal microscope was used to acquire pS129 α-syn- and TH-positive signals, as well as AOI987 and hFTAA signals, at 63× magnification.

**Whole-brain AOI987 and hFTAA** **fluorescence labeling**

The fluorescent dyes AOI987 and hFTAA were evaluated within the iDISCO+ framework in separate experiments using brain tissue from two PFF-injected mice. Each protocol followed the original procedure and included a 7-day incubation with anti-pS129 α-syn primary antibody (Abcam, ab51253, 1:1000) and anti-rabbit Cy3 secondary antibody (Jackson, 111-165-144, 1:1000). AOI987 (20 µM) or hFTAA (30 µM) was then incubated for 3 days before the tissue clearing steps. To validate the incomplete striatal labeling of α-syn aggregates and TH observed in whole-brain preparations using the original iDISCO+ protocol, cleared tissue was sectioned into 2 mm-thick coronal sections to improve antibody penetration. The sections were rehydrated through a reversed methanol series, incubated with the corresponding antibodies for 1 week at 37°C, and then recleared following the same procedure as described previously ^6^. We termed this optimized protocol “reverse” iDISCO+.

**Whole-brain immunohistochemical manual analysis of pS129** **pathology**

First, coronal brain sections with an interslice distance of 240 µm, spanning the complete rostrocaudal extent of the mouse brain, were stained for pS129 α-syn and DAPI. Next, a previously established semiquantitative grading system was used to assess the global burden of neuritic and somatic pathology ^1^. For analysis, each coronal section was matched to the corresponding ABA reference atlas image (Allen Brain Atlas, Allen Institute, USA; http://atlas.brain-map.org/), and pS129 α-syn soma and neurite pathology were graded by an experienced neuropathology expert (blinded to the grouping) in every brain region. The grading was as follows: 0 = no pS129 α-syn signal; 1 = sparse (very few neurites in the brain region or 1–3 soma in the brain region); 2 = mild (more neurites but large areas without or 4 or more soma but large areas uncovered); 3 = dense (brain region covered with neurites but spared or many soma aggregates in the brain region but places spared); and 4 = very dense (brain region densely covered with neurites or brain regions densely covered with soma aggregates). Pathology scores were assigned to specific brain regions based on the morphology of the DAPI staining and tissue autofluorescence. The mean score values were then calculated for each brain region.

**Support vector machine-based detection and local density mapping of α-syn aggregates**

For registration, the autofluorescence channel (488 nm) from the stitched LSM images was separated into fore- and background by thresholding the intensity signal and downscaling to roughly match the resolution of the GUBRA 3D-light sheet fluorescence microscopy mouse atlas ^7^. Rigid and nonrigid registration was then performed using the Elastix toolbox (affine and b-spline transform) ^8^. Nine PFF-injected mice were used to train the pipeline. First, each z-plane was median filtered using a small window (3×3 pixels) and then convolved with a Gaussian kernel (standard deviation= 1). Convolution was performed on log-transformed intensities. Next, the convolution result was thresholded (at 0.1), and aggregate candidates were detected through the identification of local maxima. The aggregates on the individual z-planes were then grouped by matching the corresponding detections on adjacent slices. This yields 3D bounding boxes, and 3D aggregate positions were defined as the centers of those bounding boxes. The aggregate candidates were then filtered by morphology using a nonlinear support vector machine (e.g., to exclude blood vessels). Analyses were performed using custom MATLAB code partially based on an open source community tool ^9^. An α-syn spread heatmap was generated by reconstructing the 3D spatial coordinates of individual aggregates from the LSM data. Each aggregate was assigned to a voxel in a 3D grid corresponding to the original image resolution. To visualize local density, we computed for each voxel the sum of aggregates within a 5-voxel neighborhood in all directions (i.e., a 5×5×5 voxel cube centered on each voxel). This procedure produces a “local density map,” highlighting regions with higher aggregate accumulation while preserving the total number of aggregates. For one brain, the heatmap was then manually aligned with the Allen Institute CCFv3 (<https://doi.org/10.6084/m9.figshare.26377171>).

**Automated quantification of TH+ quantification** **from LSM images**

For a quick assessment of TH levels and support the visual observation of dopaminergic degeneration in LSM images, we developed a complementary approach with straightforward Fiji-based quantifications. For the SNc, we defined a region of interest (ROI) on the pS129-positive MIP that encompassed most aggregates and estimated the number of TH+ cells within this ROI on three slices spaced by 60 µm. Taking advantage of the 3D dataset, we also defined two 600 × 600 × 600 µm (150 4 µm-z-plans) volumes of interest (VOIs) in the ventromedial (VOI 1) and dorsal (VOI 2) striatum and automatically quantified the TH signal intensity in the ipsilateral and contralateral hemispheres. The data were analyzed using GraphPad Prism (version 9.5.0; GraphPad Software, USA) and FiJi. All outcomes are expressed as the percentage of TH+ cells in the SNc or TH+ signal intensity in the striatum on the ipsilateral side relative to the contralateral side. Given the small sample size (N=4) and the resulting limited power to assume a normal distribution, we employed Mann‒Whitney tests, which provide a robust nonparametric alternative for comparing groups under these conditions.

**MRI data processing and analysis**

Deformation-based morphometry (DBM) was performed on all the T1W images. First, individual brains were masked using the deep learning-based lightweight framework RS2 platform to isolate brain tissue, remove nonbrain structures and manually adjust them with the software ITK-SNAP ^10^. A study-specific anatomical template was then generated through nonlinear registration, and each masked brain was aligned to this template (https://github.com/CoBrALab/optimized_antsMultivariateTemplateConstruction). The resulting deformation fields, which capture the local expansions and contractions required for alignment, were used to quantify voxelwise regional volumetric differences across the cohort to assess structural variations while maintaining anatomical correspondence to a common reference space. For statistical analysis, the Jacobian determinants representing local volumetric changes were analyzed using the FSL randomize tool. A study-specific design matrix and contrast file were created to compare groups, and a template-derived mask was applied to restrict analysis to relevant brain regions. Randomization was performed with 5000 permutations and threshold-free cluster enhancement (TFCE) to identify statistically significant voxelwise differences between groups. DTI scans underwent preprocessing, which included noise level estimation and denoising using Marchenko–Pastur principal component analysis, followed by the removal of Gibbs ringing artifacts using MRtrix3 ^11^. These steps were completed before the application of the diffusion tensor and the extraction of fractional anisotropy (FA), axial diffusivity (AD), mean diffusivity (MD), and radial diffusivity (RD) maps. The skulls were removed using the RS2 platform and manually adjusted with the software ITK-SNAP. The individual FA maps were spatially registered to the FA template from Allen Institute common coordinate framework version 3 (CCFv3) ^12^ with linear and nonlinear registration commands from AFNI software^13,^ and the generated matrices were used to register the AD, MD and RD maps. Following spatial normalization, DTI-derived metric maps were analyzed using a voxelwise permutation-based approach. FA, AD, MD, and RD maps were spatially smoothed with a small Gaussian kernel (0.3 mm full width at half maximum) within a study-specific brain mask to improve signal-to-noise while preserving anatomical specificity. For each metric, smoothed images were analyzed using the FSL randomize tool ^14^. Group comparisons were performed using predefined design matrices and contrast files modeling the four experimental groups. Statistical inference was based on 5,000 permutations and conducted using TFCE. SWI and phase images were computed using the SWI processing module in ParaVision 6.0.1 (Bruker, Ettlingen, Germany) with a Gaussian broadening of 1 mm and a mask weighting of 4. QSM aims to quantify tissue magnetic susceptibility by inverting the relationship between the measured MRI phase and the underlying susceptibility distribution. The measured phase $\phi(\boldsymbol{r})$ in a gradient-echo acquisition reflects the local magnetic field perturbation $\Delta B\left( \boldsymbol{r} \right)$, which is related to tissue susceptibility $\chi(\boldsymbol{r})$ through a convolution with the dipole kernel $d(\boldsymbol{r})$:

$$\Delta B\left( \boldsymbol{r} \right)=\gamma B_{0}\int_{V} d(\boldsymbol{r}-\boldsymbol{r}')\chi(\boldsymbol{r}')\boldsymbol{dr}'$$

where $\gamma$ is the gyromagnetic ratio, $B_{0}$ is the main magnetic field strength, and $V$ denotes the imaging volume. The inversion of this relation is inherently ill-posed because of the zeroes in the dipole kernel in k-space, necessitating specialized reconstruction techniques. In our workflow, multiecho GRE phase images were first unwrapped using a Laplacian-based phase unwrapping method ^15^ to remove phase discontinuities caused by 2π ambiguities. The background field, arising from sources outside the brain, such as air‒tissue interfaces and large veins, was subsequently removed using the V-SHARP method ^16,17^, isolating the tissue-induced phase. Finally, the local susceptibility distribution $\chi(\boldsymbol{r})$ was reconstructed from the tissue phase using the STAR-QSM algorithm ^18^, which stabilizes the ill-posed dipole inversion by thresholded k-space division and regularization, producing high-fidelity susceptibility maps with reduced streaking artifacts.

**Supplemental results**

**Evaluation of AOI987 for iDISCO+ protocol for α-syn LSM**

To evaluate whether the benzothiazole derivative AOI987 could be used with the iDISCO+ protocol and potentially serve as an alternative to the anti-pS129 α-syn antibody, we co-incubated AOI987 with the anti-pS129 α-syn antibody. While a clear α-syn pS129-positive signal was detected, AOI987 fluorescence was not visible, suggesting that the compound was likely washed out during the clearing steps (**SFig 5**).

**DAT and TH LSM comparison**

In addition, we evaluated the TH and DAT as a marker for dopaminergic cell neurodegeneration in preclinical brain clearing experiments, we applied the original iDISCO+ protocol targeting both TH and DAT in a WT mouse brain (without injection). As previously described, seven days of incubation with anti-TH primary and corresponding secondary antibodies was insufficient to achieve complete staining of the striatum (**SFig. 6a**). A comparable pattern was observed for DAT, consistent with the notion that extended antibody incubation is critical for adequate penetration in this region using the iDISCO+ protocol. Despite incomplete labeling, colocalization of TH and DAT was evident in the outer part of the striatum, suggesting that optimized conditions would likely yield full double labeling in this area, as seen in the MFB. In contrast, TH-positive cell bodies in the PVH did not exhibit detectable DAT signal. Within the SNc, DAT labeling appeared more intense around the periphery of neuronal somata and dendrites, consistent with its role as a membrane-bound transporter (**SFig. 6b**). In contrast, TH exhibited a strong signal throughout the somata and axonal projections, resulting in extensive TH/DAT colocalization, particularly within the substantia nigra reticulata (SNr). Notably, a subset of neuronal somata displayed TH expression without detectable DAT labeling, as previously reported ^19^, providing further evidence for the suitability of this 3D mapping approach and the relevance of DAT as a marker for dopaminergic circuitry.

**Dopaminergic degeneration due to α-syn injection and spreading**

To validate the quantification, we used a manual analytical approach to quantify neurons in dopaminergic nuclei in our dataset and, importantly, to study 3D TH+ signal in entire striatum. In the dorsal part of the SNc where aggregates are found, manual cell counting showed a significant reduction in the number of dopaminergic TH+ cells, consistent with Geibl *et al*., who reported an approximately 30% decrease in TH-positive neurons (**SFig 10**)^3^. Complete striatal TH staining also revealed a substantial reduction in TH-positive signal in the dorsal region of the striatum (VOI 2) (**SFig 10c, d**). These findings underscore the value of this optimized iDISCO+ protocol for achieving precise, region-specific quantification.
